# Cell states beyond transcriptomics: integrating structural organization and gene expression in hiPSC-derived cardiomyocytes

**DOI:** 10.1101/2020.05.26.081083

**Authors:** Kaytlyn A. Gerbin, Tanya Grancharova, Rory Donovan-Maiye, Melissa C. Hendershott, Jackson Brown, Stephanie Q. Dinh, Jamie L. Gehring, Matthew Hirano, Gregory R. Johnson, Aditya Nath, Angelique Nelson, Charles M. Roco, Alexander B. Rosenberg, M. Filip Sluzewski, Matheus P. Viana, Calysta Yan, Rebecca J. Zaunbrecher, Kimberly R. Cordes Metzler, Vilas Menon, Sean P. Palecek, Georg Seelig, Nathalie Gaudreault, Theo Knijnenburg, Susanne M. Rafelski, Julie A. Theriot, Ruwanthi N. Gunawardane

## Abstract

We present a quantitative co-analysis of RNA abundance and sarcomere organization in single cells and an integrated framework to predict subcellular organization states from gene expression. We used human induced pluripotent stem cell (hiPSC)-derived cardiomyocytes expressing mEGFP-tagged alpha-actinin-2 to develop quantitative image analysis tools for systematic and automated classification of subcellular organization. This captured a wide range of sarcomeric organization states within cell populations that were previously difficult to quantify. We performed RNA FISH targeting genes identified by single cell RNA sequencing to simultaneously assess the relationship between transcript abundance and structural states in single cells. Co-analysis of gene expression and sarcomeric patterns in the same cells revealed biologically meaningful correlations that could be used to predict organizational states. This study establishes a framework for multi-dimensional analysis of single cells to study the relationships between gene expression and subcellular organization and to develop a more nuanced description of cell states.

**Graphical Abstract:** Transcriptional profiling and structural classification was performed on human induced pluripotent stem cell-derived cardiomyocytes to characterize the relationship between transcript abundance and subcellular organization.

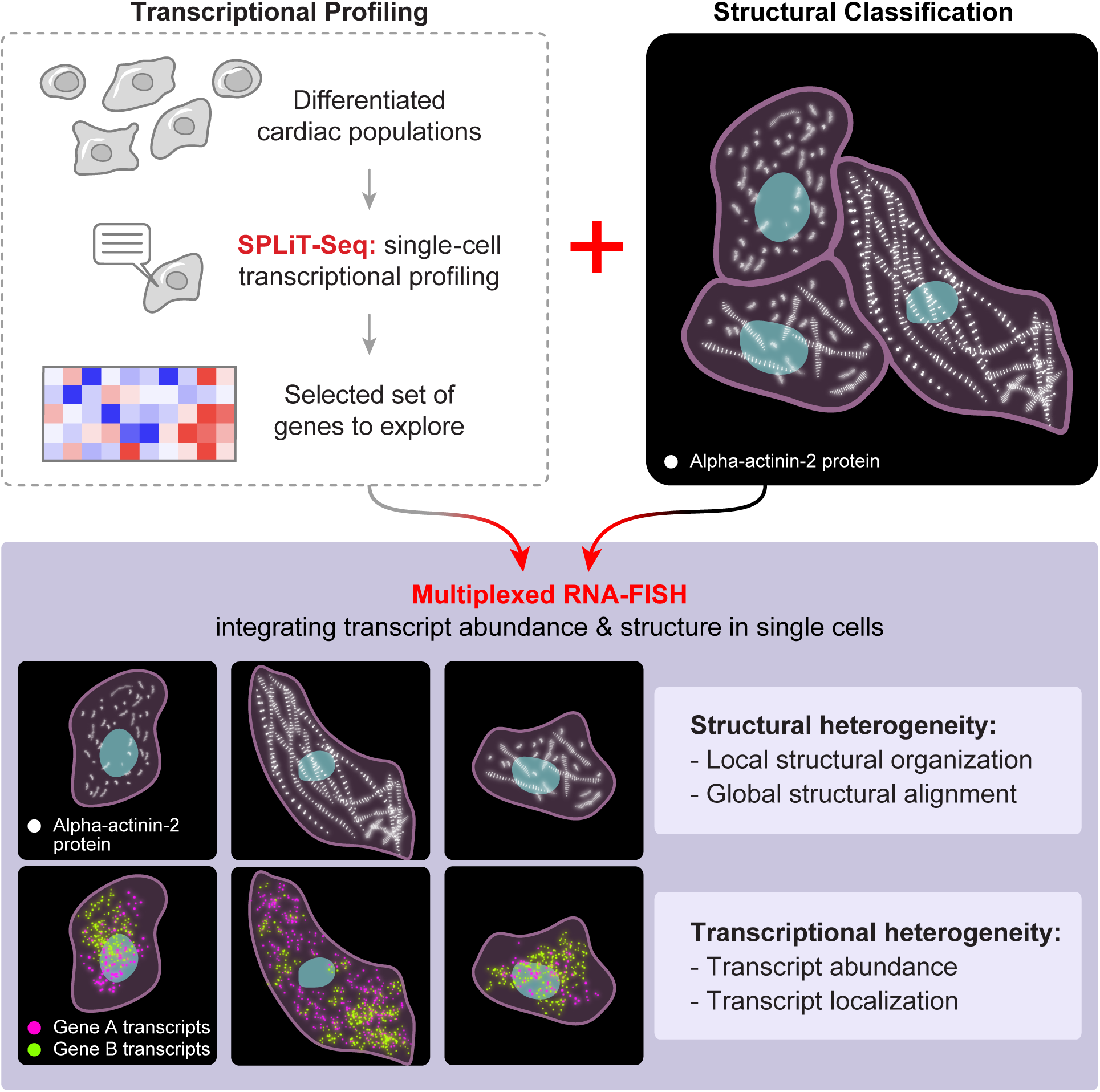

## Introduction

The earliest classifications of cells, developed well over a century ago, focused on their morphologies and locations in specific organs and tissues and revealed the intimate relationship between cell structure and function (Abbot, 1916; Ophüls, 1906). Subsequent studies enriched this relationship, showing the specialized intracellular organization of different cell types. For example, neurons have long projections that allow rapid transmission of electrical signals and synaptic specializations at their tips for cell to cell communication (Kandel et al., 2012), and muscle cells have a highly specialized contractile apparatus, permitting efficient force production (Huxley and Niedergerke, 1954; Huxley and Hanson, 1954). Even within a cell type, cells can exist in a number of functional states, e.g., the massive cellular reorganization and formation of the mitotic spindle that occurs during mitosis (reviewed in Champion et al., 2017), the polarization accompanying cell invasion (Etienne-Manneville, 2008), or the programmed cellular destruction during apoptosis (Bottone et al., 2013). The molecular era added a new dimension in cell classification with the identification of new cell types and structure-specific proteins, as well as the reorganization of generic cellular components into new specialized structures. Recently, transcriptomic data, particularly single cell RNA sequencing (scRNA-seq), added yet another dimension, revealing transcriptomic signatures of known and putative cell types during development (reviewed in Ackers-Johnson et al., 2018; Cao et al., and Paik et al., 2016; Chaudhry et al., 2019; DeLaughter et al., 2016; Kokkinopoulos et al., 2015; Lescroart et al., 2018; Li et al., 2016; 2020; Sereti et al., 2018; Suryawanshi et al., 2019; Xiao et al., 2018).

Transcriptional signatures, particularly those governed by marker genes or transcription factors driving cellular programs, are currently being widely used to identify and classify cells (Andrews and Hemberg, 2018; Shekhar and Menon, 2019). Levels of specific mRNAs in turn drive protein expression, the formation or alteration of protein complexes, and ultimately the assembly and organization of subcellular structures that perform the functions specific to those cell types. Several studies have addressed the relationship between RNA abundance and protein abundance, with Spearman (non-parametric) correlation values averaging from 0.4-0.7 (Edfors et al., 2016; Genshaft et al., 2016; Gut et al., 2018; Lundberg et al., 2010; Nusinow et al., 2020; Peterson et al., 2017; Popovic et al., 2018; Schulz et al., 2018). Ribosomal proteins, for example, show only a weak correlation between RNA expression and protein abundance due to their relative stability and highly regulated post-translational assembly into stoichiometric complexes (Schwanhausser et al., 2011). While these studies focused on linking RNA abundance to protein abundance, there is little precedent for relating RNA abundance to function (Weber et al., 2020) and organization of cellular structures (Gut et al., 2018; Popovic et al., 2018).

The goal of our study was to explore the relationship between RNA abundance and cellular organization by conjoining quantitative single cell measurements of gene expression and subcellular organization. At one extreme, it is possible that RNA abundance profiles and cell structural profiles are tightly coupled such that the cell’s structural organization (a proxy for cell phenotype) can be predicted from its gene expression, and vice versa (Liu et al., 2019; Popovic et al., 2018). However, it is also possible that known gene expression and structural classifications are connected more subtly, if at all. We hypothesized that there is a complementarity in cell organization and gene expression profiles and, therefore, combining both kinds of measurements might reveal clearer and more meaningful descriptions of cell states than either alone.

Towards this goal, we tested whether RNA abundance of key genes could predict the organizational state of the sarcomeres, the structure that powers contraction and is a hallmark of cardiomyogenesis both *in vivo* and *in vitro* (Chopra et al., 2018; Fenix et al., 2018; Sanger et al., 2005), using cardiomyocytes differentiated from human induced pluripotent stem cells (hiPSCs). The sarcomere comprises many proteins that lead to the formation of a highly regular structure that is integral to the myofibril. Differentiating cardiomyocytes show heterogeneity in sarcomeric organizational states with increasing levels of organization observed over time (Kamakura et al., 2013; Lundy et al., 2013; Snir et al., 2003). The assembly and organization of sarcomeres during cardiac differentiation occurs within the same timeframe as key transcriptional changes (Biendarra-Tiegs et al., 2019; Churko et al., 2018; Friedman et al., 2018). Therefore, we used hiPSC-derived cardiomyocytes as a model to interrogate the predictive capacity of RNA abundance with respect to structural organization on a cell by cell basis.

To quantify this relationship, we developed novel, scalable tools for the rigorous and systematic quantification of sarcomere organization in single cells and analyzed the relationship between structural organization and RNA abundance of key sarcomeric genes in the same cell. We used hiPSC-derived cardiomyocytes expressing a fluorescently tagged sarcomeric protein, alpha-actinin-2, to capture and quantify the degree of myofibril assembly. We also performed scRNA-seq to identify the gene expression signatures associated with cardiomyocyte differentiation and aging during similar timeframes. We then integrated these transcriptional and structural classifications using RNA fluorescence *in situ* hybridization (RNA FISH) (Choi et al., 2018) in these cardiomyocyte populations. This allowed us to visualize and stage structural organization quantitatively using computational algorithms, and then integrate these measurements with the expression of RNA transcript signatures in the same cell. With this approach, we systematically and robustly captured, quantified, and analyzed thousands of cells based on both structural features and RNA abundance for signature genes, creating a unique and multi-dimensional data resource for analysis.

While we found significant overlap in gene expression profiles and stages of structural organization on a population level, closer examination at single cell resolution revealed considerable heterogeneity in both gene expression and structural organization. In addition, we found a modest, but significant, correlation between the transcript abundance of *MYH7* and structural organization of the sarcomere. This myosin heavy chain is expressed in hiPSC-derived ventricular cardiomyocytes and increases in expression with maturation (Chopra et al., 2018; Yang et al., 2014). In addition, our in-depth cell-by-cell analysis showed that the organizational state of any given cell cannot be predicted from gene expression, including *MYH7*, which was the best predictor of organizational state of the population. This indicates that the measurement of both transcript abundance and subcellular organization in single cells can reveal cell states that cannot easily be identified with one measurement alone and could reveal insights that can be explored further.

In summary, we describe an innovative and scalable framework to quantify and integrate both RNA abundance and cellular organization in single cells that are challenging to measure and analyze systematically for large numbers of cells. The study highlights the power of linking single cell quantitative structural imaging with single cell transcriptional data to better understand heterogeneous cell populations. While we prototyped this concept in cardiomyocytes during the process of myofibril organization, the approach of integrating gene expression and structural organization at the single cell level can be applied more broadly to classifying cells. The integrated framework described here incorporates multiple analysis tools. These include methods for identifying transcriptomic signatures of cell states that can be used for predicting subcellular organization, quantifying structural organization at the single cell level in a way that goes beyond manual annotation or qualitative assessments, and mapping subcellular transcript localization in a systematic approach.

## Results

### Single cell RNA-sequencing of hiPSC-derived cardiomyocytes

We performed scRNA-seq to identify differentially expressed genes that define the transcriptional states and cell types present during the *in vitro* differentiation of hiPSC-derived cardiomyocytes. This included cell populations spanning four stages of the cardiac differentiation process: undifferentiated hiPSCs (Day 0), early and intermediate stage cardiomyocytes (Day 12 and Day 24), and an older cardiomyocyte population (Day 90). The early and intermediate time points (D12/24) were chosen to capture cardiomyocyte populations that are undergoing major structural and morphological changes in addition to changes in gene expression (Liu et al., 2018; Redd et al., 2019; Ronaldson-Bouchard et al., 2018). The later time point (D90) served as a benchmark for more mature cells in terms of gene expression, structure, and function (Kumar et al., 2019; Lewandowski et al., 2018; Lundy et al., 2013).

We used the combinatorial indexing-based scRNA-seq method SPLiT-Seq (Rosenberg et al., 2018), which allowed the parallel processing of all D12/D24 samples including those from two differentiation protocols, three cell lines, and multiple independent differentiation experiments (**Supplemental Table 1**). We observed similar marker gene transcript levels and cell clustering across multiple cell lines, experiments, and protocols **(Supplemental Figure 1)**. Because the transcriptomic data were consistent across protocols (**Supplemental Figure 1E, F**), all subsequent analysis was performed on experiments using a small molecule protocol (Lian et al., 2012; Lian et al., 2013) (Protocol 1, see Materials and Methods).

Unsupervised clustering of 11,619 cells collected from all four time points identified 14 clusters (Figure 1A) that segregated mainly by time point and cell type (Figure 1B, C). The cluster corresponding to undifferentiated hiPSCs (C2) was identified by the expression of the pluripotency transcription factor *POU5F1* (OCT-4) (Figure 1A-C). Five of the 14 clusters contained cardiomyocytes (C0, C1, C3, C7, C12) based on the presence of *TNNT2* transcript (Figure 1C-D), making up the majority (∼72%) of the D12/24/90 cell populations. Parallel measurement of cardiac muscle troponin T protein by flow cytometry confirmed the fraction of *TNNT2-*positive cells calculated using the scRNA-seq data (R^2^ = 0.8) (Figure 1E). The two largest clusters (C0 and C1) included a mixture of cells from both D12 and D24, with cells from D12 predominant in C0 and cells from D24 predominant in C1 (Figure 1A, 1C-bottom panel). Two other cardiomyocyte clusters (C3 and C7) represented distinct groups of D90 cardiomyocyte or cardiomyocyte-like cells, where C7 interestingly contained cells expressing cardiac (*TNNT2*), stromal (*FN1*), and smooth muscle (*TRPM3*) genes (Figure 1C). We also noted a distinct subpopulation (C12) of cardiac cells with cell cycle activity as evidenced by the expression of the proliferation marker *MKI67*, containing cells from all three differentiation time points (Figure 1C). Consistent with previous studies of cardiac populations *in vitro* (Churko et al., 2018; Friedman et al., 2018; Xu, 2012) and *in vivo* (Skelly et al., 2018), we also observed non-myocytes in the differentiated populations (∼28% of D12/D24/D90 cells). They included cells expressing markers for endothelial, stromal, smooth muscle, endodermal and ectodermal cell types (Figure 1C). These non-cardiac cell types were present in the D12/24 populations (C6, C8, C9, C11, C13) with endothelial cells (C13) being the least abundant (Figure 1C-bottom panel). Stromal, ectodermal, and smooth muscle cells were also observed in the late D90 population (C4, C5, C10); with distinct transcriptional profiles compared to D12/24 (Figure 1C). The cell types present in some clusters were challenging to define due to the presence of multiple markers associated with different cell types; C7 contained cardiomyocyte-like cells (noted above), and C5 was comprised of cells expressing stromal (*FN1*) and ECM genes (*OGN*) (Cui et al., 2019; Deckx et al., 2016) with about half of these cells also expressing the cardiac marker *TNNT2* (Figure 1C).

**Figure 1:**
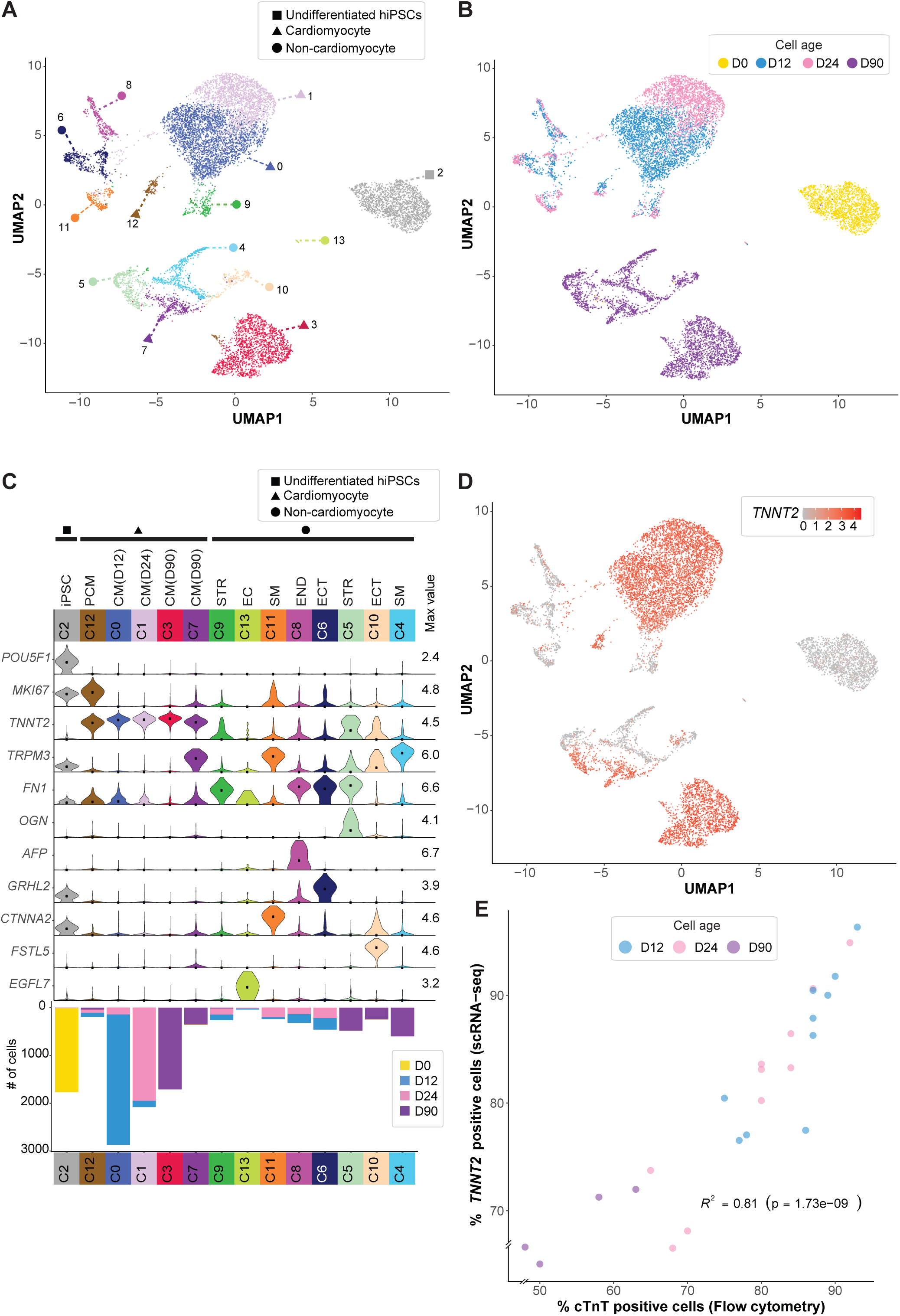
Unsupervised clustering reveals distinct cell types and cardiomyocyte clustering by time point. (A) Undifferentiated hiPSCs and hiPSC-derived cardiomyocytes (CMs) collected at 3 time points (n = 11,619) were clustered using the Jaccard-Louvain method (14 clusters indicated by colors) and visualized using UMAP. Cluster IDs were assigned after clustering based on cluster size, with C0 containing the most cells and C13 containing the least. Square icon identifies the cluster of undifferentiated hiPSCs (C2), triangles identify cardiomyocyte clusters (C0, C1, C3, C7) and the proliferative cardiomyocyte cluster (C12), and circles identify other non-myocyte cell clusters (all other clusters). (B) Same UMAP as shown in A, colored by collection time point; yellow - D0 undifferentiated hiPSCs; blue-D12 early-stage differentiated cells, pink-D24 intermediate-stage differentiated cells, purple-D90 late-stage differentiated cells. (C) Violin plots showing normalized transcript abundance of cell type marker genes by cluster (max value = maximum value of log1p normalized counts; dot = median). Bar beneath each cluster indicates cluster size (# of cells) and is colored by time point. CM = cardiomyocyte, PCM = proliferative CM, STR = stromal-like, EC = endothelial-like, SM = smooth muscle-like, END = endodermal, ECT = ectodermal. (D) UMAP colored by cardiomyocyte marker, cardiac troponin T (*TNNT2*) transcript abundance. 72% of all differentiated cells (D12, D24, D90) are *TNNT2* positive cardiomyocytes. (E) Pearson correlation of cells expressing *TNNT2* by scRNA-seq and flow cytometry: The percent of cardiac troponin T (cTnT)-positive cells from each sample determined by flow cytometry is shown in relation to the percent of *TNNT2*-positive cells from the scRNA-seq data. Individual points are colored according to time point; *R^2^* = 0.81, *p* = 1.73e^-09^.

Within the cardiomyocyte (*TNNT2* positive) populations, cells clustered along the temporal axis of differentiation: two early/intermediate cardiomyocyte clusters from D12/D24 (C0 and C1) and the late stage cardiomyocytes from D90 (C3 and C7) (Figure 1A-C). We further explored the transcriptional changes associated with these transitions by identifying the 10 genes showing the largest expression level changes (up- or down-regulated) in pairwise comparisons between the four largest clusters (C0-C3) (Figure 2A). As expected, there was a significant transcriptional shift between the hiPSC cluster (C2) and the three populations of non-proliferating cardiomyocytes (C0, C1, C3), reflecting changes in cell cycling (downregulated *MKI67*), loss of stemness (downregulated *LIN28A*, *DPPA4*) and gain of cardiac markers (upregulation of *TNNT2* and *TTN*) (Figure 2A). There were also distinct shifts in gene expression among the clusters from the early/intermediate cardiomyocyte populations (C0, C1) and the major cluster of late cardiomyocytes (C3) at D90. These included upregulation of the ventricular myosin light chain *MYL2* and the downregulation of the early/atrial myosin heavy chain *MYH6* (Bizy et al., 2013; Weber et al., 2016; Xu et al., 2009), as well as the differential expression of other structural, signaling, metabolic, and extracellular matrix (ECM) genes associated with cardiomyocyte aging and maturation (DeLaughter et al., 2016; Song et al., 2010; Xu et al., 2009; Yang et al., 2014) (Figure 2A**, Supplemental Figure 2A-C**). Of particular note was the differential regulation of several collagen genes between the clusters from D12/D24 and D90 (including an increase in *COL4A1*, *COL4A2*, *COL3A1*, *COL5A2*, and *COL19A1*, and a decrease in *COL13A1*; Figure 2A**, Supplemental Figure 2B**) that is consistent with previous observations of *in vivo* heart development and point to a remodeling of the ECM and cellular environment (Cui et al., 2018; DeLaughter et al., 2016). Finally, we also noted a decrease in *COL2A1*, a gene which has been reported to distinguish trabeculated cardiomyocytes from compact cardiomyocytes during human fetal heart development (Cui et al., 2019) (**Supplemental Figure 2B**).

**Figure 2:**
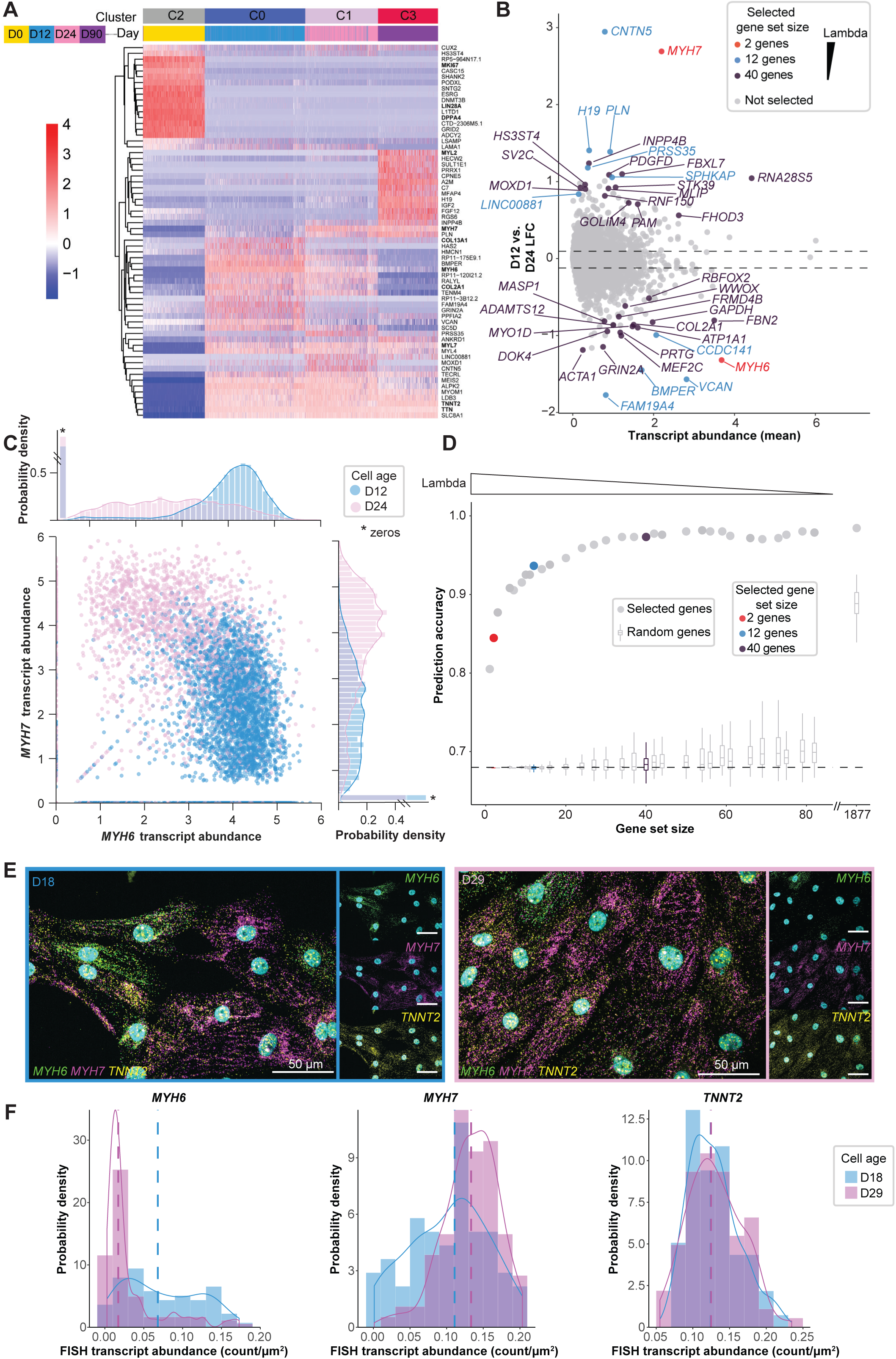
Transcriptional shifts between differentiation time points reveal differences between D12 and D24 cardiomyocytes. (A) Differentially expressed genes were identified between pairs of clusters corresponding to the undifferentiated hiPSC population (C2) and the three cardiomyocyte populations: D12 (C0), D24 (C1), and D90 (C3). Heatmap shows the top 10 up- and down-regulated genes from each pairwise cluster comparison. Normalized transcript abundance was centered and scaled across each row (color scale on left), and the dendrogram is based on hierarchical clustering of genes. Each column corresponds to a cell. Genes of interest mentioned in the text are bolded. (B) scRNA-seq mean transcript abundance (log1p of normalized counts; see Materials and Methods) vs. log2 fold change (LFC) for all genes between D12 and D24. 40 genes identified as good predictors of time point (D12 vs. D24) from bootstrapped sparse regression analysis with training data set are labeled and colored by selected gene set size; red, blue, and purple indicate gene sets selected at different values of the regularization parameter, lambda (red = 2 gene set at lambda = 0.487; blue = 12 gene set at lambda = 0.213; purple = 40 gene set at lambda 0.0494). The remaining 23,625 genes are shown in gray. The dotted lines indicate 5% and 95% quantiles of log2 fold change. (C) Scatter plot of *MYH6* vs *MYH7* normalized transcript abundance by scRNA-seq, colored by time point, with probability densities of single cell transcript abundance for each gene above (*MYH6*) and to the right (*MYH7*) of the scatter plot, normalized such that the area under each curve equals to one. Each dot represents a cell. D12 = blue, D24 = pink. *0 count cells were included in probability density calculation, but zeros were plotted separately to allow for a break in the axis. (D) Different values of the regularization parameter, lamda, were used to select gene sets, and all unique gene set sizes (x-axis) were used to calculate the prediction accuracy of cell age (dots, y-axis) in the scRNA-seq bootstrapped sparse regression test data set (see Materials and Methods). Color of selected gene set sizes are: red = 2 gene set at lambda = 0.487; blue = 12 gene set at lambda = 0.213; purple = 40 gene set at lambda 0.0494; gray = all other gene set sizes). The prediction accuracy for a set of highly variable genes (n = 1,877) is shown as the dot to the right of the x-axis break (this gene set was used for clustering and UMAP of cells shown in Supplementary Figure 2E-H). Prediction accuracies for random gene sets of the same size are shown as box plots (100 random samples) with outliers omitted. Dashed line at 0.68 indicates lower threshold for accuracy (test data set was 0.68 D12 and 0.32 D24 cells). (E) RNA FISH was performed on cardiomyocytes (differentiated from AICS-00 WTC unedited cell line) that were replated onto glass at 12 days post-differentiation and allowed to recover for 5-6 days (D18, early time point; blue), or aged an additional 2 weeks (D29, intermediate time point; pink). Transcripts are shown in green (*MYH6*), magenta (*MYH7*), and yellow (*TNNT2*), and nuclei were stained with DAPI (cyan). Representative fields of view are shown. Scale bar = 50 µm. (F) Transcript abundance for *MYH6, MYH7*, and *TNNT2* is shown as probe density (density = probe spot count/cell area in µm^2^). The colors denote time point. Histograms show probability densities, normalized such that the area under each curve is equal to one. Kernel density estimates are overlaid. (n = 70 for D18 and n = 125 for D29 cells). The dashed lines indicate median transcript abundance. See Supplemental Figure 2M for medians plotted with 95% bootstrapped confidence intervals.

We next focused on the early and intermediate time points (C0/D12, C1/D24), a developmental period comprising multiple gene expression and cellular reorganization states. Although measurably distinct from each other (Figure 1A**, Supplemental Figure 2E, F**), the changes in RNA abundance between D12 and D24 were less pronounced than those seen between the more extreme time points (Figure 2A**, Supplemental Figure 2A-D**). This is not surprising as immature cardiomyocytes undergo a program of maturation *in vivo* and *in vitro* (Cui et al., 2019; Kamakura et al., 2013; Pervolaraki et al., 2018; van Meer et al., 2016; Veerman et al., 2015). The myosin heavy chain genes *MYH6* and *MYH7* were among the relatively small set of genes with expression changes greater than two-fold (Figure 2B). When viewed at a population level, *MYH6* was more abundant at D12, and *MYH7* was more abundant at D24 (Figure 2A-C**, Supplemental Figure 2A, I-J**), an expression switch associated with the maturation of human ventricular cardiomyocytes (Bouvagnet et al., 1987; Gorza et al., 1984). However, the single cell analysis revealed a broad range of RNA abundance for *MYH6* and *MYH7* such that they were largely anti-correlated with each other with few cells expressing high levels of both transcripts (Figure 2C). Other differentially expressed genes between D12 and D24 also had broad and largely overlapping distributions with relatively modest shifts in median transcript levels between these two time points (Figure 2A**, Supplemental Figure 2A-D** and **Supplemental Table 4**).

### RNA FISH confirms population heterogeneity in gene expression

Whereas scRNA-seq provides us with a transcriptome-wide view of cardiomyocytes at the single cell level, it does not provide spatial context either at the cellular or subcellular level. Thus, we performed RNA FISH on hiPSC-derived cardiomyocytes to confirm and provide spatial context to the cell-to-cell heterogeneity in transcript abundance. Due to the technical challenges of performing RNA FISH for a large number of genes, we identified and focused on a subset of genes that most robustly distinguished the D12 and D24 populations in the scRNA-seq data. To identify these genes, we used a cluster independent bootstrapped sparse regression statistical approach (Friedman et al., 2010; Zou and Hastie, 2005). This allowed us to rank differentially expressed genes based on their ability to correctly assign individual cells to either the D12 or D24 cardiomyocyte populations (see Materials and Methods) (Figure 2B, D, **Supplemental Figure 2J-L, Supplemental Table 2**). Many of these predictive genes coincided with the differentially expressed genes between the cardiomyocyte clusters from the two time points (Figure 2A-B). Using only the expression level of the highest-ranked gene, *MYH7*, as input, a simple logistic model correctly assigned the cells to D12 or D24 with an accuracy of 0.81 (Figure 2D). Including the expression level of *MYH6*, the second ranked gene, boosted the accuracy to 0.85 (Figure 2D **– red dot)**, and including the top 12 genes raised it to 0.94 (Figure 2D **– blue dot)**. A model based on the top 40 genes performed similarly (0.97) to using all 1,877 of the most highly variable genes originally used for clustering (Figure 2D **-purple dot, Supplemental Figure 2K, L**). Therefore, a small subset of genes contained most of the relevant information that separates the transcriptional profiles of these two cardiomyocyte populations.

We first performed multiplexed RNA FISH to visualize the abundance of these transcripts in cardiomyocytes on a single cell level (three genes per cell at a time; see Materials and Methods). We initially targeted 11 differentially expressed genes including *MYH6* and *MYH7*, the two most highly predictive genes identified from the model described above (Figure 2B, E, F, **Supplemental Figure 2M** and **Supplemental Table 3**). Multiplexed RNA FISH of *MYH6*, *MYH7, and TNNT2* revealed dramatic cell-to-cell heterogeneity in expression levels, even within individual fields of view (see Materials and Methods, Figure 2E and 2F). While the absence or low abundance of certain genes in scRNA-seq data is often challenging to interpret due to potential dropouts (Kharchenko et al., 2014), the more sensitive and spatial measurement of transcripts by RNA FISH (Grün, 2014; Raj et al., 2008; Shah et al., 2016) provided independent confirmation of the cell-to-cell heterogeneity noted in the scRNA-seq data (Figure 2C, E, F).

### Cardiomyocytes show discernable stages in myofibrillar organization

The time frame when we observed cell-to-cell heterogeneity in RNA abundance of key genes by RNA FISH and scRNA-seq coincides with an analogous heterogeneity in sarcomere organization (Chopra et al., 2018; Fenix et al., 2018; Lundy et al., 2013). This offered an opportunity to test the proposition that RNA abundance levels correlate with cell organization (Battich et al., 2015; Gut et al., 2018; Popovic et al., 2018). We used an hiPSC line expressing mEGFP-tagged alpha-actinin-2 (*ACTN2-mEGFP*; Roberts et al., 2019), which localizes to the z-disk of the sarcomere, to quantify and stage sarcomere organization (Luther et al., 2011). The differentiated cardiomyocyte populations were replated for high resolution imaging and revealed a broad range of sarcomere organization with a general increase in myofibril organization over time (Figure 3A, B). Manual scoring of segmented cells by two experts (Dunn et al., 2019; Qian et al., 2013) confirmed the trend of increasing structural organization of the sarcomeres and aligned myofibrils from D18 to D32 cultures (Figure 3B, C and **Supplemental Figure 3A**). We also observed an increase in the level of mEGFP-alpha-actinin-2 protein in the more organized cells in both D18 and D32 cardiac populations (Figure 3D).

**Figure 3:**
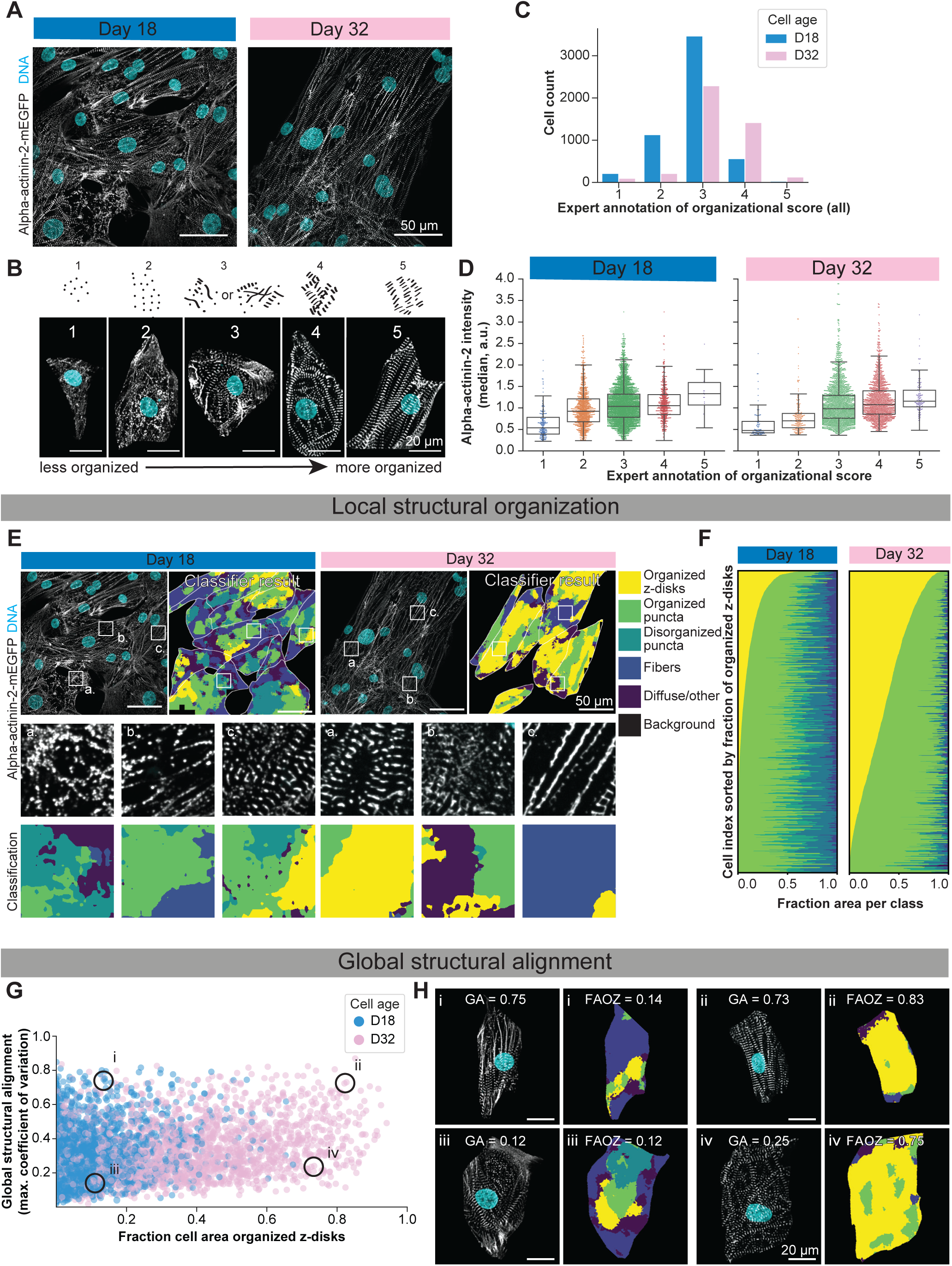
Patterns of alpha-actinin-2 organization can be classified and quantified using a deep-learning algorithm and Haralick features. (A) Representative fluorescent images of cardiomyocyte populations captured at two time points post differentiation by high-resolution spinning disk confocal microscopy. Scale bar = 50 µm. Images show heterogeneous organization of the myofibrils expressing alpha-actinin-2-mEGFP in both populations. Nuclei were stained with DAPI. Header colors indicate cell age (D18: blue, D32: pink). (B) Manual scoring of myofibril organization based on alpha-actinin-2 patterns. Variation in alpha-actinin-2 organization in individual cells were manually scored and categorized into 5 organizational classes ranging from least organized (sparse, punctate structures), score of “1”, to most organized (majority of cell area covered in regular z-disks, and well-aligned along a single axis), score of “5”. There was general concordance between the two experts (quantified in Supplemental Figure S3B). Example cells were cropped from full field of view images based on manual annotation of cell boundaries. Scale bar = 20 µm. (C) Histogram showing distribution of manually scored (expert annotation) cells into 5 myofibril organization classes. Each cell in the data set was scored by 2 expert annotators; histogram includes both scores. Colors indicate cell age (D18: blue, D32: pink). n = 4,775 cells. (D) Box and whisker plots showing the relationship between expert annotation of organizational class versus relative alpha-actinin-2-mEGFP protein intensity in arbitrary units (median a.u.) for D18 (left) and D32 cells (right). Alpha-actinin-2 protein intensity shown was calculated for each cell and divided by the median intensity for each day (D18 and D32) for a total of n = 4,775 cells. (The absolute fluorescence intensity is not comparable between D18 and D32 images due to different imaging conditions). (E) A deep learning-based classification of local structural organization of alpha-actinin-2-mEGFP. Representative fluorescent images at two time points of cardiac differentiation that show various alpha-actinin-2-mEGFP organizational patterns within a representative field of view. The original image is shown juxtaposed to the image showing the classifier results (top panel) for both populations. Header colors indicate cell age (D18: blue, D32: pink). Insets in the middle panel show examples of specific organizational features present in each population (labeled a., b., c.). The bottom row shows examples of the classifications for each inset, including organized z-disks, organized puncta, disorganized puncta, fibers, diffuse/other or background. Scale bar = 50 µm; image insets are 20 µm x 20 µm. (F) Heatmaps from two time-points in which individual cells (x-axis) have been ranked according to the fraction of the cell area consisting of organized z-disks (yellow). Colors within the heatmap represent the same organizational classes shown in panel E. n = 2,910 cells (D18), n = 2,251 cells (D32). (G) Plot showing fraction of cell area covered with organized z-disks (as in panel F) versus maximum coefficient of variation from Haralick correlation method (see “Quantification of global sarcomere alignment” in Materials and Methods for details) at D18 (blue) and D32 (pink). Cells can be separated as being regularly aligned, but containing few regular stripes, or as having high proportions of regular structure and being well aligned; examples shown in panel H of these organizational patterns are indicated with circles on the graph. n = 4,775 cells. (H) Example cells are shown from each quadrant of the graph in panel G, with alpha-actinin-2 protein (white) and nuclei (cyan). Global alignment (GA) value and fraction of area covered in organized z-disks (FOAZ) is indicated for each example cell, with color coded organizational regions representing the same organizational classes shown in panel E. Example cells were cropped from full field-of-view images based on manual annotation of cell boundaries. n = 4,775 cells. Scale bar = 20 µm.

Previous studies quantifying sarcomeric organization have relied on manual annotation of sarcomeric organization (Dunn et al., 2019) or focused on quantifying alignment and regularity of z-disk organization (Bray et al., 2008; Morris et al., 2020; Toepfer et al., 2019). With the goal of quantifying a large number of cells and achieving a finer degree of sarcomeric organizational stages, we developed a metric that captured the multiple patterns of alpha-actinin-2 associated with the sarcomere including z-disks and myofibrillar alignment (Bray, 2008; Morris, 2020; Toepfer, 2019; Pasqualini, 2015). To do so, we combined several measurements into a single weighted quantitative metric, the “combined organizational score” for automated and systematic classification of sarcomeric organization. This is an objective score that spreads out the intermediate population dominating the manual scoring (Figure 3C**, Supplemental Figure 3A**). We achieved this in three steps: 1) Developing a machine-learning based tool to measure local organization; 2) Applying an image autocorrelation method to quantify global alignment, and 3) Combining these scores together with cell morphological measurements into a linear model that not only recapitulated the expert classifications, but also provided finer quantitative discrimination between organization states.

As a first step, we devised a quantitative and scalable model for measuring alpha-actinin-2 patterns (organized z-disks, puncta, fibers, etc.) that report on local organization. We trained a pixel-based deep-learning classifier to assign regions of each cell to one of six common patterns of the alpha-actinin-2 organization, ranging from background (no visibly organized signal) to the highly regular z-disks observed for well-organized myofibrils (Figure 3E**, Supplemental Figure 3C-D**). This automated model achieved 83% accuracy in categorizing the local organization of alpha-actinin-2 in subcellular domains into each of the six pre-defined classes (**Supplemental Figure 3E-H**). We then applied this classifier across all 477 image fields (corresponding to ∼5,000 cells) in our data set. As expected, the analysis showed an increased number of cells with a regular z-disk pattern at the later time point (Figure 3F). It also confirmed and quantified the wide range of cell-to-cell variation in sarcomeric organization observed visually (Figure 3A, B), providing a finer separation of structural features than achieved by manual scoring (Figure 3F).

As a second step, we implemented a previously described global alignment measurement (Sutcliffe et al., 2018) that captured the large-scale alignment of myofibrils, an important attribute of overall structural organization of cardiomyocytes that is not captured by the local structural metric. The three global order parameters comprising this measurement describe the regularity, alignment, and spacing of the myofibrils within each individual cell independently of the total number or density of myofibrils (**Supplemental Figure 3J-K**). We identified cells with both high and low levels of both local and global organization indicating that these features of organization are unique and complementary (Figure 3G-H). One of the global alignment metrics calculated by this approach is “peak distance,” equivalent to sarcomere length. The 1.5-2 µm mEGFP-alpha-actinin-2 band spacing observed for most cells corresponds to the expected range of sarcomere length for hiPSC-derived cardiomyocytes (Lundy et al., 2013; Rodriguez et al., 2014). As expected, the median peak distance value increased slightly as the cardiomyocytes became more organized (**Supplemental Figure S3L** - lower right histogram). These shifts are not easily discernible to the human eye, highlighting the power of utilizing automated models for systematic analysis. For the final step, we combined the local organization and global organization metrics with direct measurements of cell morphology (specifically the projected cell area and aspect ratio) to generate a linear regression model (Figure 4A, B).

**Figure 4:**
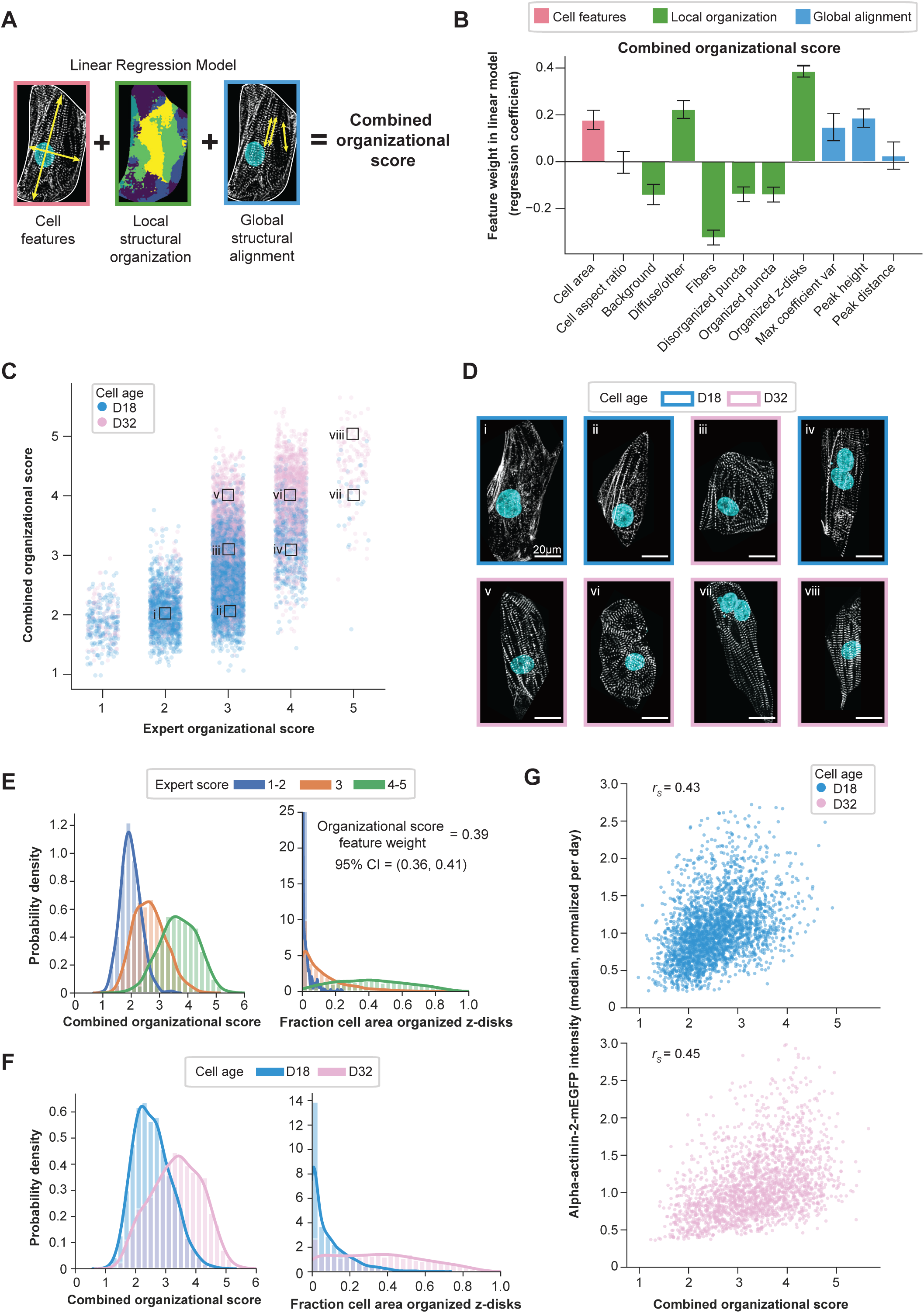
Establishing a single metric of cardiomyocyte organization by combining multiple cell features into a combined organizational score. (A) A schematic of the feature categories used in the linear regression model to compute a combined organizational score for each cell. The 3 categories represent eleven single cell metrics that capture similar features considered by experts when performing scoring: cell features (pink box) are the general cell morphological features of cell area and cell aspect ratio. Local structural organization features (green box) were derived from the automated alpha-actinin-2 classifier model and are comprised of the fraction of the cell categorized as background, diffuse/other (purple), fibers (blue), disorganized puncta (teal), organized puncta (green), and organized z-disks (yellow). Global structural alignment features (blue box) capture information about the overall alignment of the alpha-actinin-2 structure within an individual cell; here we use the maximum coefficient of variation, peak height, and peak distance (described in “Quantification of global sarcomere alignment” section of Materials and Methods). (B) Relative contribution of features to the linear regression model used to calculate a cell’s combined organizational score. The weight (regression coefficient) is plotted for each feature, colored by feature type as described in A (cell features; pink, local organization; green, global alignment; blue). Features and organizational score were calculated per segmented cell and used for further analysis. Both D18 and D32 cells were incorporated into the analysis (n = 4,775 cells in total, n = 11 total features). Error bars represent 95% bootstrapped confidence intervals (see Materials and Methods). Units for features: cell area in µm^2^; all local organization metrics (in green) as fraction of cell area; peak distance in µm. (C) Graph showing the expert organizational score per cell (x-axis) versus combined organizational score as computed using the linear regression model (y-axis). Both expert annotations are included for each cell and therefore each cell is represented twice. Each dot represents a cell, colored by time point (D18: blue, D32: pink). n = 4,775 cells. Example cells are shown in D and noted with boxes labeled i-viii. (D) Example cells from different regions of the plot representing a range of scores annotated in panel C. Alpha-actinin-2 protein (mEGFP-tagged line) shown in white and nuclei in cyan. Box boundaries indicate age of cell, D18: blue, D32: pink. Scale bars = 20 µm. (E) Histograms comparing the expert annotations of organization to the combined organizational score (left panel) and the fraction of cell area covered by z-disks (right panel), colored by expert annotation score; 1-2 (blue); 3 (orange); or 4-5 (green). Feature weight and confidence intervals are indicated. n = 4,775 cells. (F) Histograms comparing the time points with the combined organizational score calculated from the linear regression model (left), and the fraction of cell area covered by organized z-disks (right), as in panel E. Colors indicate cell age; D18: blue, D32: pink. n = 4,775 cells. (G) The combined organizational score for each cell (x-axis) plotted against the median intensity of alpha-actinin-2-mEGFP protein (y-axis-arbitrary units). Each dot is a cell, colored by cell age, D18: blue, D32: pink. Correlation coefficients (rs) for each are shown in the plot. n = 4,775 cells.

The resulting combined organizational score was generally concordant with the expert annotations (**Supplementary Figure 4A**) and also provided some interesting insights. For example, the local structural organization metric from the linear model with the strongest positive coefficient for predicting the expert annotation scores is the fraction of the cell covered by organized z-disks, and the metric with the largest negative coefficient is the fraction of area covered by fibers. This is consistent with the imaging data showing that the proportion of fibers decreased over time as the organized puncta and z-disks gained prominence (Figure 3F). We also found that cell area had a positive coefficient, suggesting increasing cell size was associated with greater cell organization, as has been reported by others (Figure 4B) (Denning et al., 2016; Karbassi et al., 2020; Snir et al., 2003).

The combined organizational score produced a nuanced and continuous quantitative metric (Figure 4C**, Supplementary Figure 4B**) that reflected the observed increase in sarcomeric organization in the D32 population as compared to D18 (Figure 4F). The fine gradation of organization offered by this combined score also achieved our goal of a more finely resolved classification of cells when compared to the expert annotation score (Figure 4E). It spread and smoothed the distribution of cells better than any individual metric, including the fraction of the cell containing organized z-disks, and resolved D32 from D18 cells more clearly (Figure 4E, F, **Supplementary Figure 4B, C**).

Since we previously noted an increase in intensity of mEGFP-tagged alpha-actinin-2 protein as cardiomyocytes became more organized (Figure 3D), we compared our combined organizational score to the relative intensity of mEGFP-tagged alpha-actinin-2 protein for each time point (Figure 4G). While the correlation is moderate for both D18 and D32 (*r_s_* = 0.43, 0.45, respectively, Figure 4G), the result is striking since it represents a direct comparison between alpha-actinin-2 protein abundance (measured by integrated intensity) and overall cellular organization for a protein that is a critical structural component of the contractile apparatus. Therefore, this measurement may serve as an upper limit for correlations between RNA transcript abundance and structural organization.

### Conjoining sarcomeric organization measurements and gene expression by RNA FISH

To integrate gene expression with structural organization in the same cells, we performed multiplexed RNA FISH targeting eight genes in cardiomyocytes expressing alpha-actinin-2-mEGFP (**Supplemental Figure 5A-B**). Four of the genes, *MYH6*, *MYH7*, *COL2A1*, and *H19*, were chosen based on the bootstrapped sparse regression analysis that identified genes best able to discriminate between D12 and D24 cardiomyocytes (Figure 2B, **Supplemental Table 2**). Three additional genes were chosen because of their role in cardiomyocyte biology: *ATP2A2*, a component of the major calcium transporter in the sarcoplasmic reticulum, *TCAP*, a component of the z-disk, and *BAG3*, a co-chaperone at the z-disk that helps maintain sarcomere function (**Supplemental Table 3**) (Judge et al., 2017; MacLennan et al., 1985; Mason et al., 1999; McDermott-Roe et al., 2019; Sakuntabhai and Timpatanapong, 1991). We selected the purine biosynthetic enzyme hypoxanthine phosphoribosyltransferase 1 (*HPRT1*) as a negative control because its abundance did not change significantly between cardiomyocyte time points in our scRNA-seq data.

A total of 4,775 manually segmented single cells from D18 and D32 cardiomyocyte populations, which were also imaged for alpha-actinin-2 structural assessment were quantified for four gene pairs using RNA FISH: *HPRT1* and *COL2A1* (n = 1,190), *MYH6* and *MYH7* (n = 1,287), *TCAP* and *BAG3* (n = 1,142), and *ATP2A2* and *H19* (n = 1,156) (**Supplementary Figure 5A-B** and **Supplementary Table 3).** Cell area correlated strongly with FISH spot counts in each cell across the transcripts evaluated (**Supplementary Figure 5C and Supplementary Table 5**), suggesting that transcript count may be regulated at the level of mRNA density or concentration rather than absolute number (Battich et al., 2015; Padovan-Merhar et al., 2015; Tempe et al., 2015; Torre et al., 2018). To decouple this relationship, we calculated FISH spot densities (count/µm^2^) for all eight genes and found a broad range of absolute expression levels and significant cell-to-cell heterogeneity in both D18 and D32 cell populations (**Supplementary Figure 5B)**.

To determine whether RNA abundance correlates with cell structural organization, we evaluated the transcript abundance with respect to the combined organizational score on a cell-by-cell basis for all eight genes (Figure 5A). To decouple changes in RNA abundance that were only associated with cell age from changes associated with cell organizational state, we calculated the Spearman rank correlation coefficient (*r_s_*) for each of the D18 and D32 cell populations separately as well as for the combined cell population (Figure 5B). *HPRT1*, the negative control, showed a modest but statistically significant negative correlation (*r_s_* = −0.21) for the combined population (Figure 5B). While this establishes a floor for a biologically meaningful correlation between RNA abundance and structural organization, the correlation of 0.43-0.45 between protein abundance of alpha-actinin-2 and structural organization set the upper limit (D18 and D32, respectively, Figure 4G, Figure 5B).

**Figure 5:**
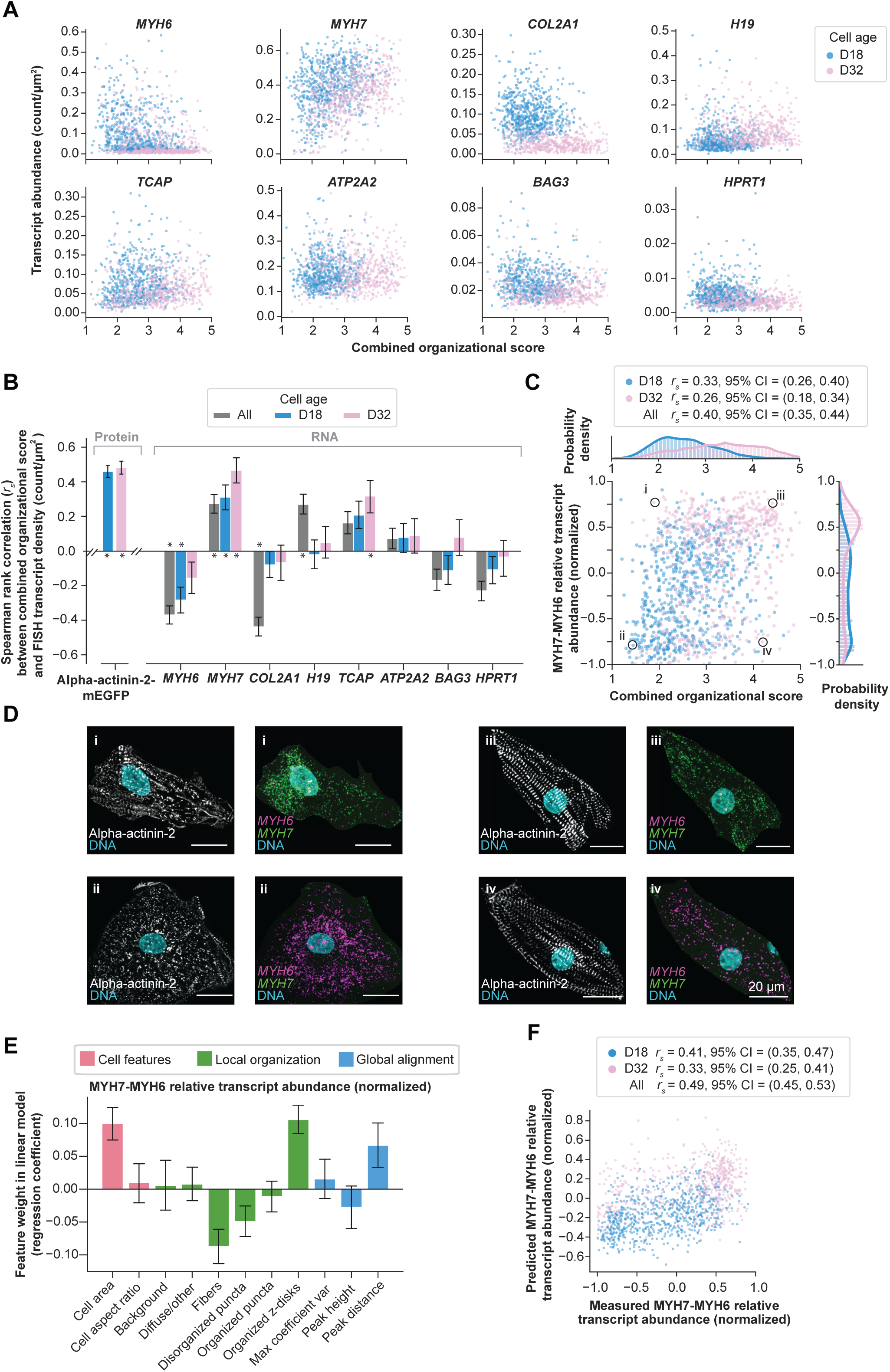
Analysis of structural organization and gene expression reveals correlations. (A) Scatter plots showing transcript abundance from RNA FISH, reported as density (count/µm^2^) (y-axis) versus combined organizational score (x-axis) for each evaluated gene. Transcript abundance is measured by FISH probe spot density (counts per µm^2^) in single cells. Each plot represents one gene and each dot represents a single cell (n = 1,287 cells for *MYH6* and *MYH7*, n = 1,190 cells for *COL2A1* and *HPRT1*, n = 1,156 cells for *H19* and *ATP2A2*, n = 1,142 cells for *BAG3* and *TCAP*). Colors indicate cell age (D18: blue, D32: pink). (B) Bar plots showing the Spearman rank correlation between combined organizational score and either alpha-actinin-2 protein abundance (mEGFP intensity) or transcript abundance of the genes listed. Correlation metrics are shown for each of the eight genes that were probed by RNA FISH in alpha-actinin-2-mEGFP cardiomyocytes for each time point and as combined populations. The correlation between alpha-actinin-2 protein abundance with combined organizational score establishes our upper limit for expected biological correlation, while the correlation of the control gene *HPRT1* establishes the lower limit. Grey bars indicate the correlation across all cells; blue and pink bars show the correlation for cells stratified by age (D18: blue, D32: pink) and asterisks indicate correlations with biological significance. Error bars are 95% bootstrapped confidence intervals. Measurements where the bootstrapped interval crosses zero are not statistically significant. (C) Scatter plot with normalized relative abundance of *MYH7* compared to *MYH6* transcripts (y-axis) calculated from RNA FISH transcript abundance plotted against the combined organizational score (x-axis) across single cells (n = 1,287) (see Materials and Methods for details of relative expression calculation; levels of *MYH7* were 4-5x higher than those of *MYH6*, thus the difference between these two genes is scaled from −1 (100% *MYH6*) to +1 (100% *MYH7*), with the median of 0 representing cells expressing both genes. Colors indicate cell age (D18: blue, D32: pink). Top histogram shows the marginal distribution of the organizational scores stratified by cell age as a probability density normalized such that the area under each curve is equal to one. Right histogram shows the marginal distribution of the relative expression of *MYH7* and *MYH6*, also stratified by cell age. Spearman correlations (*r_s_*) and confidence intervals (CIs) are as noted in the figure with “All” referring to correlations of combined D18 and D32 cells. Circles marked by Roman numerals (i-iv) refer to example cells shown in panel D. (D) Example images of four cells representing a range of transcript abundance and combined organizational scores as indicated in panel C, showing alpha-actinin-2-mEGFP protein (white) on the left for each cell and *MYH6* transcript (magenta), and *MYH7* transcript (green) by RNA FISH on the right. DNA is shown in cyan. Example cells were cropped from full field-of-view images based on manual annotation of cell boundaries. Scale bar = 20 µm. (E) Relative contribution of cell features to the linear regression model used to calculate a cell’s predicted *MYH7* vs *MYH6* relative transcript abundance. The feature weight (regression coefficient, y-axis) is plotted for each feature, colored by feature type (cell features: pink, local organization: green, global alignment: blue). Features were calculated for each segmented cell and used for further analysis. Both D18 and D32 cells were incorporated into the analysis, n = 715 cells (D18) and n = 572 cells (D32). Error bars indicate 95% bootstrapped confidence intervals. (F) Scatter plot of *MYH7* and *MYH6* relative transcript abundance (density difference, as shown in panel C) showing predicted values calculated from linear regression model shown in panel E (y-axis) and measured values (x-axis). Spearman correlations (*r_s_*) and 95% confidence intervals (CI) are indicated in the legend with “All” referring to correlations of combined D18 and D32 cells. Each dot represents a cell, colors indicate cell age (D18: blue, D32: pink). n = 715 cells (D18) and n = 572 cells (D32).

*ATP2A2* and *BAG3* showed no significant correlation between RNA abundance and combined organizational score at either time point or in the combined population compared to negative control (*r_s_* = 0.07 and −0.16 for the combined cell population, respectively, Figure 5B). We observed an interesting pattern of correlation for *COL2A1* and *H19*, both of which increased in RNA abundance over time; but this increase was not related to the combined organizational score. Specifically, the correlation within either the D18 or D32 populations for both were negligible (*r_s_* < −0.08 and < 0.05, respectively), whereas the correlation in the combined population was strongly negative for *COL2A1* (*r_s_* = −0.4) and modestly positive for *H19* (*r_s_* = 0.25, Figure 5B). The change in average RNA abundance for both these genes as measured by RNA FISH between D18 and D32 is readily apparent (Figure 5A**, Supplemental Figure 5B**) and the direction of the abundance change (decreasing over time for *COL2A1* and increasing over time for *H19*) agrees with the scRNA-seq data for these two genes (Figure 2B). However, there is no significant correlation between RNA abundance and combined organizational score within either D18 or D32 populations for both genes (Figure 5B), despite exhibiting a great deal of structural and gene expression heterogeneity. Therefore, this change in RNA abundance appears to be strictly time-dependent and is not correlated with the increase in cellular organization (Figure 4F). This result underlines the importance and the direct utility of measuring both cellular organization and RNA abundance jointly on a cell-by-cell basis, rather than relying on separate measurements.

Three genes, *MYH6*, *MYH7* and *TCAP*, exhibited meaningful correlations with the combined organizational score for at least one of the single-day populations (Figure 5B). RNA abundance for *MYH7* showed significant correlations with organization for both time points independently, with the largest correlation observed for D32 (*r_s_* = 0.43). *TCAP* also showed a positive correlation with the strongest at D32 (*r_s_* = 0.29). In contrast, *MHY6* negatively correlated with organization (*r_s_* = −0.34 in the combined cell population, Figure 5B), which is consistent with the decrease observed in RNA abundance over time between D18 and D32 (Figure 5A, Supplemental Figure 5B). The path from RNA expression to sarcomere assembly encompasses many layers of regulation and the highly coordinated assembly of dozens of distinct proteins and protein complexes into the contractile sarcomere. Therefore, it is remarkable that this degree of correlation between transcript and organization can be directly observed on a cell-by-cell level within the heterogeneous population of intermediate-stage cardiomyocytes. Intriguingly, the level of correlation observed for *MYH7* and organization (*r_s_* = 0.43) is consistent with the correlation between protein abundance of alpha-actinin-2 and the combined organizational score (Figure 4G **and** Figure 5B), suggesting that this level of correlation is highly biologically significant.

We further examined the relationship between cell structural organization and the relative expression levels of the two myosin heavy chain genes *MYH6* and *MYH7* (Figure 5C), genes whose expression switches during maturation of early ventricular cardiomyocytes (Racca et al., 2015; Reiser et al., 2001; Xu et al., 2009). Because RNA abundance of *MYH7* correlated positively and *MYH6* correlated negatively with the combined organizational score, we evaluated whether the ratio of expression would provide a more powerful prediction of cell organizational state. We found that the measurement of the relative abundance of RNA for these two alternative myosin heavy chains did not enhance the predictive value with respect to the combined organizational score as compared to the abundance of *MYH7* alone (correlation *r_s_* =0.33 and 0.26 for D18 and D32, respectively, Figure 5C). Therefore, these results show a correlation between *MYH7* abundance and cell organization despite the high level of cell-to-cell heterogeneity. The single cell analysis also revealed individual cells with high levels of both sarcomeric organization and relative RNA abundance of *MYH6*, and, conversely, cells with low levels of organization but high relative abundance of the *MYH7* transcript (Figure 5C, D). This suggests that the gene expression switch where *MYH6* is largely replaced with *MYH7* does not cause the increase in structural organization of the myofibril observed during this time frame. It also highlighted the importance of the cell-by-cell evaluation of both gene expression and structural organization compared to population analysis.

These results demonstrate that the combined organizational score, developed to refine the quantitation of expert annotations, shows a meaningful correlation between RNA abundance and cell structural organization. Next, we attempted to directly predict gene expression (Figure 5E) using the same 11 features as the previous model for predicting cell organization (Figure 4B). We used the relative *MYH7-MYH6* RNA abundance as the target for the prediction to retain the maximal amount of informative gene expression signal (Figure 5C). The Spearman correlation between the *MYH7-MYH6* relative transcript abundance as predicted by this simple linear model and the actual measured values was *r_s_* = 0.41 and 0.33 for D18 and D32, respectively (Figure 5F). These values were comparable to the correlation with the combined organizational score that relied on expert annotation for its development. This striking result indicates that it is indeed possible to predict gene expression (on a statistical, population level) from image-based cell organizational features, using only the top pair of correlated genes in our study. The fraction of the cell covered by organized z-disks and fibers produced the strongest positive and negative coefficients, respectively, in this model as well as in the previously described model for combined organization score (Figure 4B, Figure 5E). This provides strong circumstantial evidence that features useful for predicting the patterns recognized by the experts are similar to the image-based cellular features connecting sarcomere organization to relative transcript abundance of *MYH7-MYH6*. Overall, these analyses revealed a broad range of organizational features that could be exploited to understand the relationship between gene expression and structural organization in an automated fashion.

### Transcripts reside in distinct subcellular locations

RNA FISH also revealed a variety of distinct patterns of subcellular transcript localization within these cardiomyocyte populations (reviewed in Buxbaum et al., 2015; Park et al., 2012; Popovic et al., 2018). Among the eight transcripts evaluated, the *H19, BAG3, TCAP and HPRT1* transcripts were primarily distributed throughout the cytoplasm while *COL2A1* and *ATP2A2* displayed perinuclear localization (Figure 6A **and Supplemental Figure 6B**). These classes of the cytoplasmic and perinuclear transcript localization were confirmed quantitatively by measuring the relative distance between each FISH spot and the cell nucleus, as normalized to the nearest position of the cell plasma membrane (Figure 6B **and Supplemental Figure 6C**). In contrast, *MYH7* and *MYH6* localized to the myofibrils in a subset of cells (Figure 6A **and Supplemental Figure 6A**), which was not captured by these measurements. Therefore, we annotated sarcomeric localization of these transcripts on a cell-by-cell basis manually. Interestingly, this annotation of over 700 cells showed the *MYH7* transcript co-localized with myofibrils in >60% of the cells (Figure 6D). This sarcomeric localization of *MYH7* was slightly more prevalent in the cells from D32 as compared to D18, although not significant (Figure 6D). In contrast, it was rare to observe this sarcomeric localization pattern in cells expressing *MYH6* transcript (<10% of cells), potentially due to its low level of RNA abundance observed in these cells (Figure 6C). While sarcomeric localization of transcripts has been reported in rat myocytes (Kehat et al., 2011; Lewis et al., 2018), to our knowledge, this has not been previously described in human stem cell-derived cardiomyocytes.

**Figure 6:**
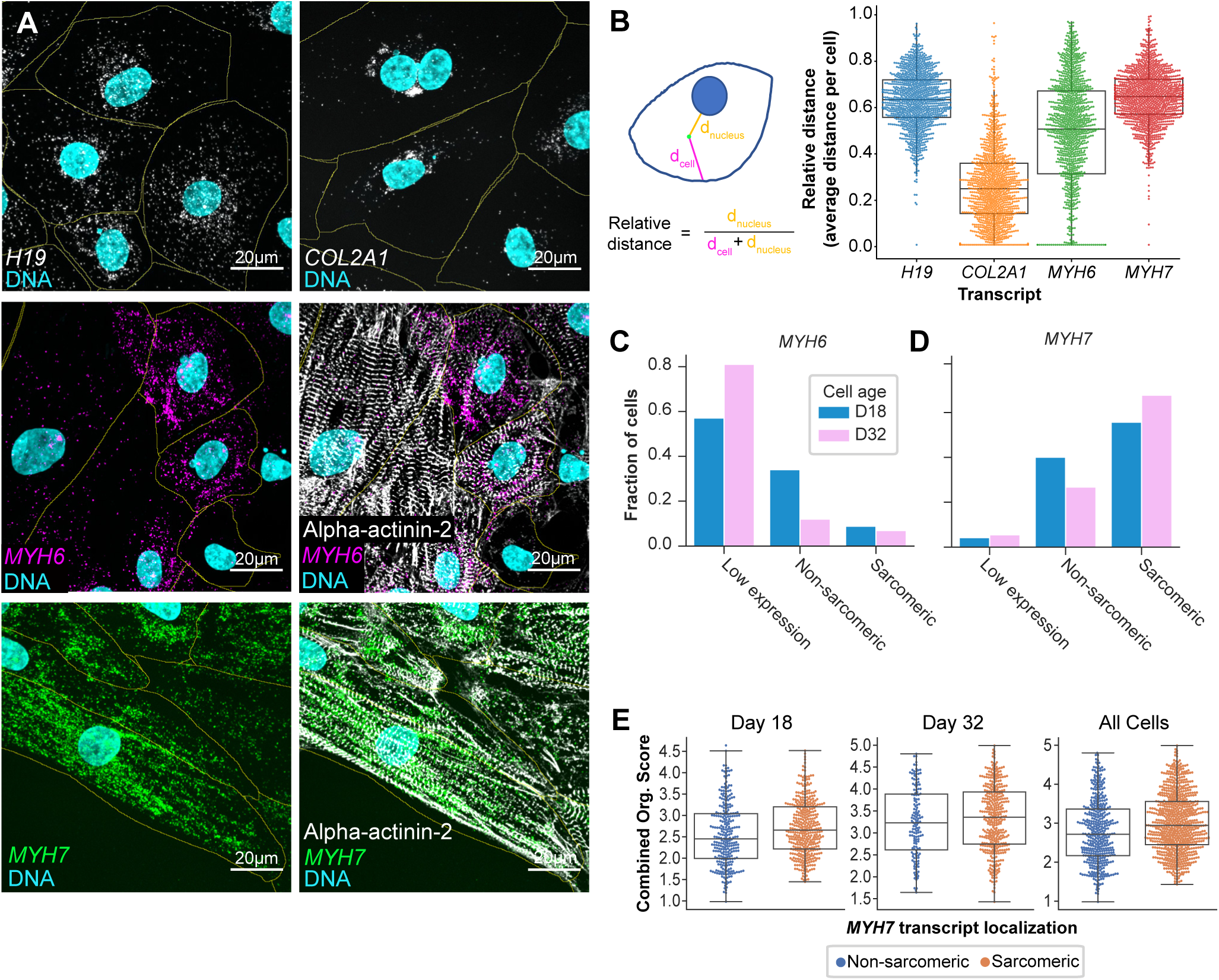
Transcripts display distinct subcellular localization patterns. (A) Representative images of cytoplasmic (*H19*, white), perinuclear (*COL2A1*, white), and myofibrillar (*MYH6*, magenta and *MYH7*, green) subcellular transcript distribution in cardiomyocytes. In each image, yellow lines indicate manually-drawn cell boundaries for clarification and nuclei are shown in cyan. In *MYH6* and *MYH7* images (middle and bottom rows), the right panel shows alpha-actinin-2 protein (endogenously tagged with mEGFP, white). Scale bars are 20 µm. Images were cropped to show more detail of individual cells. (B) The schematic (left panel) of the method used to evaluate localization patterns. The relative distance between transcript and nucleus was calculated for each gene studied by RNA FISH, where d_n = distance from nucleus and d_c = distance from cell boundary. The right panel shows these values for *H19*, *COL2A1*, *MYH7*, and *MYH6*. Each data point represents the average relative distance of the transcript in a single cell. Combined data for D18 and D32 time points are shown for all four transcripts. n = 1,287 cells for each *MYH6* and *MYH7*, n = 1,190 cells for *COL2A1*, n = 1,156 cells for *H19*. (C) Manual annotation was performed to categorize the localization of *MYH6* transcripts within each cell at D18 (blue) and D32 (pink) as: low expression, non-sarcomeric, or sarcomeric. Fraction of total cells evaluated per time point are shown for each category. n = 713 cells (D18) and n = 564 cells (D32). (D) Manual annotation was performed to categorize the localization of *MYH7* transcript at D18 (blue) and D32 (pink). Distribution was recorded as in panel C. n = 713 cells (D18) and n = 564 cells (D32). (E) Swarm plots showing the distribution of cells with sarcomeric and non-sarcomeric *MYH7* transcript localization (x-axis), both per day and in aggregate, plotted against the combined organizational score for each cell (y-axis). Left panel shows cells from D18 (n = 683 cells), middle panel shows cells from D32 (n = 533 cells), and right panel shows cells from both days combined (n = 1,216 cells).

The correlation between *MYH7* RNA abundance and overall cell structural organization raised the question whether the localization of *MYH7* transcripts to the myofibrils is more strongly associated with its organizational state. We examined the distribution of combined organizational scores for cells with either sarcomeric or non-sarcomeric localization of *MYH7* (Figure 6E) and did not observe a statistically significant trend (Figure 6E). We also tested this computationally by calculating the relative densities of *MYH7* transcripts in regions of cells associated with organized z-disks (**Supplemental Figure 6E**). We first determined the degree of enrichment of transcript localization in these areas over a random distribution of probes, then compared this enrichment score to the combined organizational score (**Supplemental Figure 6F**). While this measurement does not directly capture the qualitative manual annotation, this analysis also showed neither a correlation between subcellular localization of *MYH7* transcript and cell structural organization, nor a strong relationship between sarcomeric localization of *MYH7* transcript and its overall abundance (**Supplemental Figure 6D-F**).

## Discussion

In this study, we established a proof-of-concept platform, combining quantitative image-based readouts of gene expression, transcript localization, and cellular organization. We developed and used this multi-dimensional analysis to determine the feasibility of predicting the organizational state of a cell from its transcriptional state and vice versa, with the ultimate goal of developing better methods to classify cell states than could be achieved by either transcriptomic measurements or cellular morphological measurements alone.

We prototyped this approach by evaluating both transcriptional and structural changes in hiPSC-derived cardiomyocytes during the early/intermediate stages of differentiation, a period in which there are changes in both gene expression and myofibrillar organization of the contractile apparatus. We used scRNA-seq to identify transcriptional changes during this process, exploiting the variation in gene expression levels comprising the heterogeneous cardiomyocyte populations, and focused on a few key genes that are associated with myofibril assembly and organization for co-analysis with structure. To identify and stage structural organization in these cardiomyocytes, we developed a “combined organization score” based on a set of image-based metrics including machine learning classifiers that captured, quantified, and parameterized organizational states of the sarcomere in each cell. We then used multiplexed RNA FISH, along with localization of mEGFP-tagged sarcomeric alpha-actinin-2 protein, to capture and quantify a range of gene expression, localization, and structural changes in the same cell population and study these relationships cell-by-cell.

We found that the RNA abundance of a single gene, *MYH7*, best predicted the degree of myofibrillar organization. This is consistent with the switch from *MYH6* to *MYH7* expression being a key discriminator between cardiomyocyte populations collected during the time frame of increasing myofibrillar organization in our studies and others (Pioner et al., 2016; Racca et al., 2015; Reiser et al., 2001; Xu et al., 2009; Yang et al., 2014). Furthermore, *MYH6* and *MYH7* were the top pair of genes identified from our statistical model as the best predictors of cell stage based on scRNA-seq data. Although the correlation between *MYH7* abundance and sarcomeric organization was relatively modest at ∼0.4, it is biologically significant as it reflects the relationship between transcript abundance and a highly intricate structure (the sarcomere) and is similar to the correlation between alpha-actinin-2-mEGFP protein intensity and sarcomeric organization. This is especially striking considering the direct correlation between RNA and protein is generally only in the range of 0.4-0.7 (Edfors et al., 2016; Genshaft et al., 2016; Gut et al., 2018; Lundberg et al., 2010; Nusinow et al., 2020; Peterson et al., 2017; Popovic et al., 2018; Schulz et al., 2018).

This systematic single cell co-analysis of RNA abundance and structural organization also revealed a wide range of sarcomeric organization in cells expressing either or both *MYH6* and *MYH7.* Intriguingly, we found well-organized cells expressing high *MYH6* and low *MYH7*, and poorly organized cells with high *MYH7* and low levels of *MYH6*. This suggests that the presence of the *MYH7* transcript, although generally correlated with high levels of organization, is not required to produce or maintain an organized sarcomere and that the expression of *MYH6* alone is compatible with the presence of organized sarcomeres. These observations are consistent with a recent study showing low correlations between the expression of myosin proteins and cellular electrophysiology (Weber et al., 2020).

The image-based co-analysis of transcript and structure also revealed interesting subcellular transcript distribution patterns including the localization of *MYH7* transcripts along the sarcomere. While sarcomeric localization of *Myh6*, *Myh7* and *Actc1* transcripts has been previously reported in neonatal and adult rat cardiomyocytes (Buxbaum et al., 2015; Kehat et al., 2011; Lewis et al., 2018), this is the first report of their localization in differentiating hiPSC-derived cardiomyocytes, suggesting that localization of sarcomere protein-encoding transcripts may also play a role in myofibril assembly and organization in human cells. While we did not find an obvious relationship between transcript localization and organization of the sarcomere in this study, additional analysis of sarcomere-associated transcripts with structural imaging may provide insights into the role of transcript localization patterns in myofibril organization and maintenance. Furthermore, incorporating subcellular landmarks with RNA FISH can be used to construct comprehensive maps of transcript localization that go beyond sarcomerogenesis.

Taken together, our findings suggest that cell states may be better classified by a combination of spatial and quantitative measurements of transcripts and structural analysis rather than by either one alone. While gene expression profiles capture the stages during which myofibril assembly occurs and can parse it into early and late stages, they do not easily capture the many, readily categorized structural states that comprise the process of myofibril organization we observed in our cell-by-cell analysis nor the localization of transcripts to specific subcellular regions. This complicates the simple notion of a state-to-state transition and adds nuance to predicting cell states from gene expression profiles and vice versa. Additional measurements, such as functional phenotype (force generation, electrophysiology, and metabolism for cardiomyocytes) might further sharpen the cell categorization. Finally, by combining these measures with larger subsets of genes using highly multiplexed FISH or *in situ* sequencing [e.g., MERFISH (Chen et al., 2015; Wang et al., 2018); seqFISH (Eng et al., 2019), or FISSEQ (Lee et al., 2015)], even more granular and meaningful classifications may emerge. In this context, the present study is only a first step towards building an understanding of the relationships that could be used to predict cellular phenotypes.

Previous related studies have described the use of RNA FISH in tissues to retain spatial, morphological, and neighborhood information that is generally lost during cell dissociation for single cell RNA-sequencing (Battich et al., 2015; Gut et al., 2018; Lopez et al., 2019; Popovic et al., 2018; Torre et al., 2018). This enabled both the visualization and spatial characterization of different cell types within intact three dimensional tissues (Moffitt et al., 2016; Shah et al., 2016) and interrogation of the relationship between cell type, neighborhood, and gene expression in tissue culture (Battich et al., 2015; Gut et al., 2018; Popovic et al., 2018; Raj et al., 2008). Our study extended this general approach to cellular organization, in our particular case the formation and organization of myofibrils, describing a set of tools to classify states of subcellular organization in a highly quantitative, systematic, and automated fashion. These metrics go beyond the current set of tools available for subcellular analysis of myofibril organization in single cells (Bray et al., 2008; Dunn et al., 2019; Morris et al., 2020; Toepfer et al., 2019) and enable integration and correlation of transcript measurements with structural organization. This framework of quantifying and merging multiple measurements can be applied to many other data types with an ultimate goal of predicting phenotype from observable data including structural organization.

In summary, we have developed a novel framework for combining single cell gene expression with quantitative measurements of cellular structure that can be scaled for large numbers of cells and additional measurements. We developed machine learning-based image quantitation and classification tools and used various statistical approaches to select a subset of state-defining genes from large scRNA-seq data sets and predict cellular features. By leveraging these tools, we identified gene expression and structural states and demonstrated the feasibility and limits of predicting cell organization from RNA abundance and vice versa. Although broad transcriptomic signatures may be sufficient for classifying major cell types such as the epithelial-like pluripotent stem cells versus mature cardiomyocytes versus fibroblasts, deciphering more nuanced differences in cell types, or the states that they occupy will benefit from the kind of integrated approach presented here. While the specific image-based tools we developed are specific for myofibril organization, analogous methodologies can be developed for other structures and cell types. For example, ion channel isoform shifts during cardiomyocyte maturation will likely best be measured with transcriptomics or electrophysiology, whereas the enormous structural rearrangements in the intermediate stages of differentiation seem more readily classified by image-based analysis. This study represents a step towards the goal of conjoining high dimensional single cell data derived from large cell populations to create a new and holistic understanding of cell behavior.

## Acknowledgements

We thank Joy Arakaki for flow cytometry analysis, Joyce Tang for RNA FISH image collection, Colette DeLizzo and Haseeb Malik for cardiomyocyte maintenance and sample collection, Derek Thirstrup for CellProfiler guidance, Thao Do for illustrations, and Jianxu Chen for technical discussions. We would like to thank the Allen Institute for Cell Science founder, Paul G. Allen, for his vision, encouragement, and support.

## Author Contributions

Conceptualization, K.A.G, T.G., M.C.H., R.D.M., S.R.J.T., and R.N.G.; Methodology, R.D.M., M.P.V., C.Y., K.A.G., T.G., M.H., M.F.S. and S.R.; Software, J.B., G.R.J., M.P.V., M.F.S., C.Y., R.D.M., and T.G.; Formal Analysis, T.G., R.D.M., M.P.V., M.F.S. and C.Y.; Investigation, K.A.G., T.G., M.C.H., M.H., A.N., C.M.R., A.B.R., C.Y., R.J.Z., S.Q.D., A.N., and J.L.G.; Data Curation, R.D.M., T.G., J.B., C.Y., and M.P.V.; Writing – Original Draft, K.A.G., T.G., M.C.H., and R.D.M.; Writing – Review and Editing, R.N.G., J.A.T., S.M.R. K.A.G., T.G., M.C.H., R.D.M., and K.R.C.M.; Visualization, K.M., K.A.G., T.G., M.H., R.D.M., R.J.Z., and M.P.V.; Supervision, R.N.G., J.A.T., S.M.R., G.S., N.G., T.K., V.M., and S.P; Project Administration, K.R.C.M., R.N.G., and K.A.G., M.C.H.

---

**Supplemental Figure 1, related to** Figure 1**: scRNA-seq cell transcriptional profiles of cells from the early (D12/14) and intermediate (D24/26) time points from multiple differentiation experiments**

A. Early and intermediate time point (D12/14 and D24/D26) cells differentiated with either Protocol 1 (small molecule, D12/24 cells in Figure 1) or Protocol 2 (cytokine, D14/26 cells) representing three cell lines and five independent differentiation experiments were clustered together using the Jaccard-Louvain method. Cells were visualized using UMAP and colored by cluster (17 clusters).
B. Same UMAP shown in panel A, colored by collection time point; Protocol 1 (small molecule) = D12 and D24. Protocol 2 (cytokine) = D14 and D26.
C. Same UMAP shown in panel A, colored by *TNNT2* transcript abundance. 78% of cells are *TNNT2* positive cardiomyocytes.
D. Violin plots showing distributions of transcript abundance for cell type marker genes by cluster from A (max value = maximum value of transcript abundance; dot = median). Bar beneath each cluster indicates cluster size (# of cells) and is colored by time point (top panel), differentiation experiment (middle panel), and cell line (bottom panel).
E. Spearman rank correlation coefficient was calculated using aggregate counts for each gene between the two protocols at proximate time points, D12 (Protocol 1) vs D14 (Protocol 2). Each dot represents one gene. Raw counts for each gene were summed across all cells from each time point and log1p-transformed.
F. Same analysis as shown in panel E, but for D24 (Protocol 1) vs D26 (Protocol 2).
G. Spearman correlations for global transcriptional profiles between each pair of cell lines. Each dot represents a gene.
H. Spearman correlations for global transcriptional profiles between each pair of differentiation experiments. Each dot is a gene.
I. Density kernel plots showing the transcript abundance distributions of cardiac marker genes for cells from each protocol at proximate time points, D12 (blue) vs D14 (gray), and D24 (pink) vs D26 (purple); *TNNT2* = cardiac muscle troponin T, *MYH6* = myosin heavy chain 6, *MYH7* = myosin heavy chain 7, *MYL7* = myosin light chain 7, *MYL2* = myosin light chain 2. Y-axis is probability density.

**Supplemental Figure 2, related to** Figure 1 **and** Figure 2**: scRNA-seq transcriptional shifts between cardiomyocyte time points with focus on D12 and D24**

A. Transcript abundance distributions for structural genes with temporal transcriptional shifts between D0 (C2), D12 (C0) cardiomyocytes, D24 (C1) cardiomyocytes, and D90 (C3) cardiomyocytes (small molecule/protocol 1; see Figure 1, 2). The dots within the violin plots indicate median transcript abundance for each gene. Max denotes maximum transcript abundance and the asterisks indicate genes from the top 40 selected genes that best distinguish D12 and D24 cells in the bootstrapped sparse regression analysis (see Figure 2B, Supplemental Table 2).
B. Same as A, but showing ECM related collagen genes.
C. Same as A and B, but showing genes involved in signaling and metabolic shifts.
D. Transcript abundance distributions for other genes that are differentially expressed between D12 and D24; same cells and time points as in panel A. As in panels A-C, asterisks indicate genes from the top 40 selected genes that best distinguish D12 and D24 cells in bootstrapped sparse regression analysis (see Figure 2B, Supplemental Table 2).
E. Cells from C0 (D12 cardiomyocytes) and C1 (D24 cardiomyocytes) in Figure 1A were clustered independently of other cells. UMAP colored by cluster. Clusters are numbered i0-i5 (i is used to distinguish these clusters from those shown in Figure 1).
F. Same UMAP shown in panel E colored by time point.
G. Same UMAP shown in panel E colored by differentiation experiment.
H. Same UMAP shown in panel E colored by *TNNT2* (cardiac troponin T) transcript abundance.
I. Transcript abundance distributions for marker genes in clusters i0-i5 (UMAP of clusters shown in E). i0, i3, i4 are predominantly D12 cells; i1 and i2 are predominantly D24 cells; i5 is a mix of D12 and D24 cells.
J. Heatmap of 40 genes that best distinguish D12 and D24 cells in bootstrapped sparse regression analysis shown in clusters i0-i5, which are comprised of D12 and D24 cardiomyocytes (see Figure 2B, Supplemental Table 2).
K. Frequency of selection for genes in 1,000 bootstrap rounds of logistic binomial regression with the elastic net penalty. In each bootstrap round, 100 models were fit using a sequence of 100 lambda values (the regularization parameter). X-axis shows log of lambda, and y-axis shows number of times gene (max 1,000) had a non-zero coefficient for a given value of lambda. Dotted line indicates value of lambda used to define selection (0.0494). All genes are shown. Genes in gray were not in the set of 40 selected genes (see Figure 2B, Supplemental Table 2). The 40 selected genes are further colored to show the two top 2 selected genes (red), the top 12 selected genes (blue), and the rest of the selected genes from the set of 40 (purple).
L. Number of selected genes at each value of the regularization parameter, lambda. The 40, 12, and 2 gene sets are colored as in K and Figure 2B, D.
M. Median RNA FISH transcript density (count/µm^2^) is shown for 11 genes in AICS-00 WTC (unedited, non-structure) cardiomyocytes (see Supplemental Table 3) with bootstrapped (n = 1,000) 95% confidence intervals. Number of cells per gene *COL2A1*: D18 = 169, D30 = 157; *ESRRG*: D18 = 349, D30 = 291; *FABP3*: D18 = 148, D30 = 117; *H19*: D18 = 148, D30 = 117; *MEIS2*: D18 = 372, D30 = 283; *MYH6*: D18 = 418, D30 = 382; *MYH7*: D18 = 418, D30 = 382; *MYL2*: D18 = 687, D30 = 595; *MYL7*: D18 = 539, D30 = 478; *TNNT2*: D18 = 405, D30 = 443; *VCAN*: D18 = 169, D30 = 157.

**Supplemental Figure 3, related to** Figure 3**: Classification of alpha-actinin-2 using ML-based and Haralick features**

A. Additional examples showing diversity of cells that were scored as a “3” (see Figure 3B, C); examples were cropped from full field-of-view images based on manual annotation of cell boundaries. Because annotations were assigned based on the majority of cell area, cells with widely varying levels of organization could all fall into the “3” bin. Scale bar = 20 µm. Alpha-actinin-2 protein-mEGFP (white), DNA (cyan).
B. Confusion matrix showing concordance between the two expert annotators in scoring alpha-actinin-2 organization in single cells from combined D18 and D32 populations. n = 4,775 cells.
C. Examples of five distinct patterns of alpha-actinin-2 organization identified by a human annotator and scored as shown in Fig 2C; these correspond to organized z-disks (wide, regularly oriented bars, yellow), organized puncta (dots with a clear axis of organization, green), disorganized puncta (dots without a clear axis of orientation, teal), fibers (long, thread-like patterns, blue) and diffuse/other patterns (diffuse or overlapping areas with no clear organizational pattern, purple). Representative example annotations are shown as insets from the whole field-of-view. Scale bar = 50 µm, image insets are 20 µm x 20 µm.
D. Results of the deep-learning based classification of local structure organization in the whole field-of-view and sub-regions shown in panel C. Scale bar = 50 µm, image insets are 20 µm x 20 µm. Colors represent organizational class, as in panel C.
E. Improvement in deep learning classifier model accuracy over training epochs.
F. Model loss across training epochs. Loss refers to the function that is used to determine how accurate the model currently is and to calculate gradients for optimization. A lower loss score is indicative of a more accurate model.
G. Confusion matrix showing concordance between human class annotations and model predictions.
H. Evaluation of model confidence in predictions. The model outputs a five element vector; each element has a value between 0 and 1, representing the probability (or amount of “confidence” the model has) that the image patch is of the corresponding class, with all elements summing to 1. The purple line shows the model’s accuracy when restricted to a given confidence threshold; the blue line shows the percent of classifications in the validation set which had a confidence above a given value.
I. Comparison of live and fixed image showing the classification model, which was trained on data collected with live cardiomyocytes, transfers well to fixed samples. Example shown is a field-of-view that was imaged live, and the same position was imaged after fixation. Insets show features and classifications using the same classification groups and color scheme shown in panel C. Scale bar = 50 µm, image insets are 20 µm x 20 µm.
J. Metrics for an example cell with a high fraction of regular stripes and low global alignment. Image shows alpha-actinin-2 protein (white). Top plot shows offset distance versus Haralick correlation for varying rotational angles (solid lines) and exponential fitting of curves (dashed lines). Bold black line highlights the curve for rotational angle with maximum correlation. Middle plot shows offset distance versus the correlation of exponential fitting; the peak distance and peak height metrics are indicated with green arrows. Bottom plot shows offset distance versus the mean, coefficient of variation and standard deviation of the Haralick correlation; the maximum coefficient of variation is indicated with a green arrow. See Materials and Methods for a complete description of calculations.
K. As in panel J, metrics for an example cell with a high fraction of regular stripes and high global alignment. Image shows alpha-actinin-2-mEGFP protein (white).
L. Plots showing correlations between features calculated to measure global alignment (Max coefficient var, Peak height, and Peak distance) and expert annotation score (blue/orange/green) or cell age (pink and blue). n = 4,775 cells.

**Supplemental Figure 4, related to** Figure 4**: Expert annotator agreement and linear model input**

A. Confusion matrix panels showing concordance of the combined organizational score (rounded to the nearest whole number) with the expert annotation classification that was summed from two annotators (left) or individual expert annotation (middle and right). Diagonal represents agreement from expert annotation and the combined organizational model. n = 4,775 cells scored by each expert.
B. Violin plot colored by cell age, D18: blue, D32: pink, showing the spread and increase in the organization scores from manual expert annotation (x-axis) versus combined organizational score as computed using the linear regression model (y-axis). Data is the same as what is shown in Figure 4C, n = 4,775 cells.
C. Histograms of each of the features used as input for the linear model used to derive the combined organizational score are shown in relation to expert annotation of organization or cell age. Features are organize by type: cell features (enclosed by pink box), local organization (enclosed by green box), or global alignment metric (enclosed by blue box); histograms colored by expert annotation score (top) given as 1-2 (orange), 3 (blue), or 4-5 (green), or colored by cell age (bottom), D18: blue, D32: pink. Organizational score feature weight and confidence interval is displayed for each feature (also shown in Figure 4B).

**Supplemental Figure 5, related to** Figure 5**: Transcript abundance and localization varies by gene and scales with cell area**

A. Representative images of cells after performing RNA FISH for *MYH6*, *MYH7*, *COL2A1*, *HPRT1*, *BAG3*, *TCAP*, *H19* and *ATP2A2*. Imaging shows alpha-actinin-2-mEGFP protein (white), DNA (cyan), and transcripts (magenta or green). Example cells were cropped from full field-of-view images based on manual annotation of cell boundaries. Scale bar = 20 µm.
B. Distribution of transcript abundance for each gene is shown as transcript count (left) or density in count/µm^2^ (right). Colors indicate cell age (D18: blue, D32: pink).
C. Scatter plots showing transcript abundance as transcript count (y-axis) versus cell area in µm^2^ (x-axis) for each transcript. Each dot represents the total transcript count for each gene in a single cell (n = 1,190 cells for *HPRT1* and *COL2A1*, n = 1,287 cells for *MYH6* and *MYH7*, n = 1,142 cells for *TCAP* and *BAG3* and n = 1,156 cells for *ATP2A2* and *H19*). Colors indicate cell age and corresponding Spearman correlations (D18: blue, D32: pink, All; gray) where “All” refers to the correlation for the combined D18 and D32 cell populations. Confidence intervals are listed in Supplementary table 5.

**Supplemental Figure 6, related to** Figure 6**: Transcripts display distinct subcellular localization patterns**

A. Example images of transcript localization of *MYH7* showing both sarcomeric and non-sarcomeric (non-sarc) RNA localization. In all images, *MYH6* transcript is magenta, *MYH7* transcript is green, alpha-actinin-2-mEGFP protein is white and DNA is cyan. Scale bars are 20 µm. White boxes indicate regions enlarged for detail on the right. Images have been cropped for clarity.
B. Representative images of spatial distribution of *ATP2A2*, *BAG3*, *HPRT1*, and *TCAP*. In each image, transcripts are shown in white, nuclei are shown in cyan, and manually-drawn cell borders are shown in yellow. Scale bars are 20 µm. Images have been cropped to show more detail of individual cells.
C. Relative distance of each transcript from the nucleus from RNA FISH images was computed using the same method as described in Figure 6B. Each data point represents the average relative distance of the transcript in a single cell. Combined data from both time points are shown for each gene. n = 1,156 cells for *ATP2A2*, n = 1,142 cells for *BAG3* and *TCAP*, n = 1,190 cells for *HPRT1*.
D. Histogram of *MYH7* transcript density in cells manually classified as having non-sarcomeric (blue) and sarcomeric (orange) localization of *MYH7* transcript. n = 1,287 for both time points; n = 715 cells for D18 and n = 572 cells for D32.
E. Histogram of enrichment of *MYH7* transcripts in regions of cells associated with organized mEGFP-tagged alpha-actinin-2 z-disks (as identified by the deep-learning classifier). This was done in cells manually annotated as having sarcomeric (orange) or non-sarcomeric (blue) localization of *MYH7.* n = 1,287 cells for both time points; n = 715 cells for D18 and n = 572 cells for D32.
F. Combined organizational score of a cell plotted against the enrichment of *MYH7* transcript in regions of the cell categorized as having organized z-disks, organized puncta, fibers, and all of those categories combined (n = 1,287 cells). Colors indicate cell age (D18: blue, D32: pink).

**Supplemental Table 1: scRNA-seq sample metadata**

Metadata for all samples included in the scRNA-seq data set. Sequencing_batch refers to the sequencing batch that samples belonged to (seq1 or seq2), cell_line indicates one of three cell lines: either AICS0 (AICS-00 WTC), AICS11 (AICS-0011 cl.27 TOMM20-mEGFP), or AICS37 (AICS-0037 cl.172 TNNI1-mEGFP). Protocols are listed as small molecule (referred to as Protocol 1), small molecule (7.5/7.5) (also Protocol 1 with an optimized small molecule concentration; described in detail in the “Directed Cardiomyocyte Differentiation” section of the Materials and Methods), and cytokine (Protocol 2). Differentation_experiment refers to the experiment batch, Exp 1 through Exp7, which corresponds to the differentiation_start. Differentiation_start refers to the date at which undifferentiated stem cells were seeded for cardiac differentiation, with the undifferentiated cell passage number and seeding density noted in 10×10^6 cells per well (M, million) in columns H and I, respectively. The harvest date and day when spontaneous beating was observed is recorded (# of days after D0, when differentiation was initiated, see Directed cardiomyocyte differentiation section of Materials and Methods, nr = not recorded). Percent_ctnt is the percent of the harvested population that expressed cardiac troponin T by flow cytometry analysis.

**Supplemental Table 2: Feature selection genes ranked by lambda**

40 selected genes (see Figure 2B, 2D, and Supplemental Figure 2K, 2L, and scRNA-seq feature selection analysis section of Materials and Methods) from feature selection analysis ranked by lambda value (regularization parameter) when they were first selected. Selected = gene with non-zero coefficient in all 1,000 bootstrap rounds.

**Supplemental Table 3: Genes evaluated using RNA FISH**

Genes evaluated using RNA FISH were chosen in four categories: 1) feature selection analysis between D12 to D24 (described in Figure 2B, 2D, and Supplemental Figure 2K, 2L, scRNA-seq feature selection analysis section of Materials and Methods, and Supplemental Table 2), 2) differential expression with a prominent change in transcript abundance in scRNA-seq analysis from D24 to D90, 3) known biological relevance in cardiomyocyte differentiation or sarcomere organization, and 4) control (category for each gene listed in Column F). RNA FISH probe sets for each listed gene can be ordered from Molecular Instruments/Molecular Technologies using the unique probe ID listed here (Column D). Protein name (Column B) and NCBI accession number (Column C) for the sequence used to design probe sets is also given. RNA FISH was performed either in AICS-00 WTC (unedited) or AICS-075 ACTN2-mEGFP cardiomyocytes as described in relevant text and figure legend and as listed in Column H. RNA FISH data set from AICS-00 (unedited) cells is referred to as “non-structure” because of the lack of fluorescent sarcomere structure channel, and data set from AICS-075 ACTN2-mEGFP cells is referred to as “structure” because of presence of sarcomere structure channel.

**Supplemental Table 4: Differentially expressed genes between clusters**

List of differentially expressed (DE) genes from pairwise cluster comparisons for D0 (C2), D12 (C0), D24 (C1), and D90 (C3) (see clusters in Figure 1A and scRNA-seq differential expression analysis section of Materials and Methods). Each tab is DE genes between one pair of clusters. LogFC = log2 fold change between groups; logCPM = mean log2 of counts per million; LR = likelihood ratio statistics; PValue = p-value before multiple testing correction; FDR = multiple testing adjusted p-values with Benjamini-Hochberg method to control false discovery rate; up = fraction of non-zero cells for gene in up-regulated cluster (cluster in pair with higher transcript abundance of the two)r; down = fraction of non-zero cells for gene in down-regulated cluster (cluster in pair with lower transcript abundance).

**Supplemental Table 5: Spearman correlation values and confidence intervals for cell area versus transcript abundance**

List of Spearman correlation values for plots shown in Supplemental Figure 5C, which show cell area in µm^2^ (x-axis) versus transcript abundance (y-axis). Each row specifies the Spearman correlation value and 95% confidence interval for genes in D18, D32 and All populations, where “All” refers to the correlation for the combined D18 and D32 cell populations. Number of cells included in calculation are provided in the Supplemental Figure 5C legend.

## MATERIALS AND METHODS

### RESOURCE AVAILABILITY

#### Lead contact and materials availability

Further information and requests for resources and reagents should be directed to and will be fulfilled by the Lead Contact, Ru Gunawardane.

### MATERIALS AVAILABILITY

This study did not generate new reagents or cell lines. All gene-tagged hiPSC lines used in this study can be obtained through the Allen Cell Catalog (www.allencell.org/cell-catalog) and the unedited WTC-11 line from Coriell (Coriell Institute GM25256).

### DATA AND CODE AVAILABILITY

Data are available in the Allen Cell open resource data package accessible at: https://open.quiltdata.com/b/allencell/tree/aics/integrated_transcriptomics_structural_organization_hipsc_ cm/

Analyses were performed using the code deposited in the Allen Cell Modeling github repository accessible at: https://github.com/AllenCellModeling/fish_morphology_code

Manual annotation was performed using Napari, accessible at: https://github.com/Napari/napari, https://github.com/AllenCellModeling/napari-annotation-tools

## EXPERIMENTAL MODEL AND SUBJECT DETAILS

### Cell Lines

All work with human induced pluripotent stem cell (hiPSC) lines was approved by internal oversight committees and was conducted in accordance with NIH, NAS, and ISSCR guidelines. The male WTC-11 (WTC) hiPSC line was generated and generously provided by Dr. Bruce Conklin (The Gladstone Institute) (Kreitzer et al., 2013) and referred to as AICS-00 in the manuscript. Genetically edited hiPSC lines used in this study were generated as previously described (Roberts et al., 2017; Roberts et al., 2019). These include AICS-0011 cl.27 (TOMM20-mEGFP), AICS-0037 cl.172 (TNNI1-mEGFP), and AICS-0075 cl.85 (ACTN2-mEGFP), and can be obtained through the Allen Cell Collection (www.allencell.org/cell-catalog; Allen Institute for Cell Science, 2020).

## EXPERIMENTAL METHOD DETAILS

### Human induced pluripotent stem cell culture

All hiPSC lines were maintained in mTeSR1 (Stem Cell Technologies #85850) supplemented with 1% penicillin/streptomycin (P/S) (Gibco #15070063) on plates coated with growth factor reduced matrigel (Corning #354230). hiPSCs were passaged as single cells after reaching 70-85% confluency every 3-4 days using Accutase (Gibco #A11105-01) and replated in mTeSR1 supplemented with 1% P/S and 10 µM Rock Inhibitor (Y-27632, Stem Cell Technologies #72308). A detailed, step by step protocol is available at the Allen Cell Explorer (www.allencell.org/methods-for-cells-in-the-lab, SOP: WTC culture v1.7.pdf) (Allen Institute for Cell Science, 2020).

### Directed cardiomyocyte differentiation

For single cell RNA-sequencing experiments, two differentiation protocols were used, referred to as “Protocol 1” and “Protocol 2” (**Supplemental Figure 1, Supplemental Table 1)**. Protocol 1 is based on a previously reported small molecule differentiation method with modifications (Lian et al., 2012; Lian et al., 2013). Briefly, hiPSCs were dissociated into a single cell suspension using Accutase and seeded into matrigel-coated 6 well-plates at 125k-350k cells per well in mTeSR1 supplemented with 1% P/S and 10 µM Rock Inhibitor (denoted as Day −3). Cells were fed daily with mTeSR1 media for 2 days, and on the third day (Day 0) differentiation was initiated with the addition of RPMI-1640 (Invitrogen #A10491-01) supplemented with B27 without insulin (Invitrogen #A1895601) and 6 µM CHIR99021 (Cayman Chemical #13122). At 48 hours (Day 2) media was replaced with RPMI-1640 supplemented with B27 without insulin and 5 µM IWP2 (R&D Systems #3533). After an additional 48 hours (Day 4), media was replaced with RPMI-1640 supplemented with B27-without insulin, and replaced every 2-3 days thereafter (starting on Day 6) with RPMI-1640 supplemented with B27 containing insulin (Invitrogen #12587010) and 1% P/S. All cardiomyocytes generated for Day 90 samples and for RNA FISH experiments were differentiated using an optimized version of Protocol 1 in which Chiron and IWP were both used at 7.5 µM (Roberts et al., 2019). A detailed, step by step protocol is available at the Allen Cell Explorer (www.allencell.org/methods-for-cells-in-the-lab<u>, SOP:</u> Cardiomyocyte differentiation methods_v1.2.pdf) (Allen Institute for Cell Science, 2020).

Protocol 2 is a previously reported method using a combination of cytokines and small molecules to induce cardiac differentiation (Palpant et al., 2015). Briefly, hiPSCs were dissociated into a single cell suspension using Accutase and seeded into matrigel-coated 6 well-plates at 1×10^6^ - 2×10^6^ cells per well in mTeSR1 supplemented with 1% P/S, 10 µM Rock inhibitor, and 1 µM CHIR99021 (denoted as Day-1). Differentiation was initiated the following day (Day 0) with the addition of RPMI-1640 supplemented with B27 without insulin, 100 ng/mL ActivinA (R&D Systems #338-AC), and 1:60 diluted growth factor reduced Matrigel. After 17 hours (Day 1), media was replaced with RPMI supplemented with B27 without insulin and containing 1 µM CHIR99021 and 5 ng/ml BMP4 (R&D systems #314-BP). After an additional 48 hours (Day 3), media was replaced with RPMI-1640 supplemented with B27 without insulin and 1 µM XAV 939 (Tocris Biosciences #3748). After an additional 48 hours (Day 5), media was again replaced with RPMI-1640 supplemented with B27 without insulin, and cultures were fed with RPMI-1640 supplemented with B27 containing insulin and 1% P/S every 2–3 days thereafter (starting on Day 7).

Across both differentiation protocols, spontaneous beating was generally observed between Days 7-12.

Please note that we have recently updated our protocols and recommend the use of RPMI-1640 (Invitrogen #11875-093) and B27 supplement (Gibco #17504044) for cardiac differentiation using both protocols.

### Experimental design for scRNA-seq

For each differentiation experiment, one source plate of hiPSCs was dissociated and seeded concurrently for both differentiation protocols, thus keeping the input hiPSCs the same. Samples were collected on the same day for each paired collection time point listed in the scRNA-seq data set, from both protocols (Day 12/14, and Day 24/26, **Supplemental Table 1**). The difference in reported day (i.e. Day 12 vs 14) is due to the delay in differentiation initiation (Day 0) between the two protocols, as described above in the “Directed cardiomyocyte differentiation” section of the Materials and Methods. Samples were collected 15 days after seeding (denoted as D12 for Protocol 1, D14 for Protocol 2) or 27 days later (denoted as D24 for Protocol 1, D26 for Protocol 2). Samples referred to as D90 were independently derived and were collected 93-96 days after initiating differentiation using a modified version of Protocol 1, as described above in the “Directed cardiomyocyte differentiation” section of the Materials and Methods. Each scRNA-seq sample represents a single well, and experiment replicates were independently derived (see **Supplemental Table 1**). Stem cell (D0) samples for scRNA-seq were independently cultured and did not precede the differentiation setups. All scRNA-seq sample metadata can be found in **Supplemental Table 1**.

### Single cell dissociation for scRNA-seq

Stem cells were dissociated to single cells as described above in the “Human induced pluripotent stem cell culture” section of the Materials and Methods and processed for RNA sequencing following the protocol detailed in the “scRNA-seq library preparation and sequencing” section in the Methods below. For differentiated cardiomyocyte samples, differentiated wells were visually inspected at the desired cardiomyocyte collection time point for successful cardiac differentiation on the basis of high cardiac purity, spontaneous beating, and morphology; wells that passed these criteria were collected for downstream analysis. To dissociate cardiomyocytes into single cells, wells were first washed with 1X DPBS (Gibco #14190-144) and incubated with pre-warmed 2X TrypLE Select (Gibco #A12177) diluted in Versene (Gibco #15040-066) for 8-10 minutes at 37°C. Monolayers were gently dissociated into a single cell suspension using a P1000 micropipette and added to 5 mL of RPMI-1640 containing B27 supplement, 1% P/S, 10 µM Rock Inhibitor and 200 U/mL DNAse I (Millipore Sigma #260913-10MU) (resuspension media). Wells were washed twice with an additional 1 mL of resuspension media, and the cell suspension was centrifuged at 300 g for 5 minutes at 4°C. Cells were gently resuspended in 5 mL of resuspension media and counted twice on a hemocytometer to obtain a total cell count for each sample.

### Flow cytometry of scRNA-seq samples

At the time of single cell dissociation as described above, a minimal aliquot was taken from each cell sample for cardiac Troponin T (cTnT) analysis by flow cytometry as previously described (Roberts et al., 2017). Briefly, aliquots were fixed in 4% Paraformaldehyde (Electron Microscopy Sciences #15710) in DPBS for 10 min. Fixed samples were stained in BD Perm/Wash^TM^ buffer (BD Biosciences #512091KZ) containing anti-cardiac Troponin T AlexaFluor® 647 (BD Biosciences #565744) or equal mass of AF647 lgG1 κ isotype control (BD Biosciences #565571) for 30 min followed by resuspension in 5% FBS (Gibco #10437028) in DPBS with 2 µg/mL DAPI before processing and data acquisition on a CytoFLEX S (Beckman Coulter). Analysis was conducted using FlowJo software V. 10.2 (Treestar).

### scRNA-seq library preparation and sequencing

The remainder of the cells from the single cell dissociation described above in the “Single cell dissociation for scRNA-seq” section were prepared for SPLiT-seq library preparation by first centrifuging the remaining cell suspension at 300 g for 5 minutes at 4°C and then resuspending in 1 mL of cold RNAse-free PBS containing 0.05 U/µL Superase IN (Invitrogen #AM2696) and 0.05 U/µL Enzymatics RNAse Inhibitor (Qiagen #Y9240L) (mixture referred to throughout as PBS + RI). On ice, resuspended cells were passed through a 40 µm filter and fixed with 3mL of 1.33% formaldehyde Electron Microscopy Sciences #15710) for 10 minutes. Fixed cells were permeabilized with 160 µL of 5% Triton X-100 (MilliporeSigma #T8787-50ML)+ RNase Inhibitor for 3 minutes. Permeabilized cells were then centrifuged at 500 g for 5 minutes at 4°C and resuspended in 500 µL of PBS + RI, and mixed with an additional 500 µL of 100 mM Tris-HCL (ThermoFisher #AM9855G) and then 20 µL 5% Triton X-100. Cells were then centrifuged at 500 g for 5 minutes at 4°C and the pellet resuspended in 300 µL 0.5X PBS + RI then passed through a 40 µm filter. Filtered cells were counted using a hemocytometer and diluted to 1×10^6^ cells/mL in 0.5X PBS + RI. Labeled tubes were placed in a RT (room temperature) Mr Frosty (Thermo Fisher #5100-0001) and placed into a −80°C Freezer for storage.

In-cell reverse transcription, ligation barcoding, lysis, and library preparation were carried out according to the version 3.0 protocol at https://sites.google.com/uw.edu/splitseq and described in (Rosenberg et al., 2018). Vials were thawed by placing tubes in a 37°C water bath, and vial contents were pipetted into wells containing barcoded, well-specific reverse transcription primers and a reverse transcription reaction mix. Each well contained a mixture of random hexamer and anchored poly(dT)_15 barcoded RT primers. Two different sequencing experiments were conducted. Sequencing experiment 1 contained samples across Day 12/14 and Day 24/26, while sequencing experiment 2 contained samples across D0-Stem Cells, Day 90, and one re-sequenced sample each of Day 12 and Day 24 that had been included in experiment 1 to control for sequencing batch variability. Sample metadata including sequencing experiments can be found in **Supplemental Table 1**. For sequencing batch 1 (48 samples; see **Supplemental Table 1**), each sample was placed in a single well (4,000 cells per input sample) for a total of 192,000 cells. For sequencing batch 2 (9 samples, see **Supplemental Table 1**), each D0 and D90 sample was split over 6 wells (24,000 input cells per sample). The D12 and D24 samples in sequencing batch 2 were each split into 3 wells (12,000 input cells per sample). In both sequencing batches, after three rounds of barcoding the cells were counted and divided into sub-libraries of 5000 cells before lysis. These sub-libraries were barcoded with a fourth unique barcode each and processed for sequencing on an Illumina NextSeq.

### scRNA-seq data processing

Sequence alignment and quantification of intronic and exonic UMI (unique molecular identifier) counts were performed as described in (Rosenberg et al., 2018) using STAR (Dobin et al., 2013) for alignment, TagReadWithGeneExon from Drop-seq tools (Macosko et al., 2015) to assign reads to genes, and Starcode (Zorita et al., 2015) to collapse UMIs. Per gene intronic and exonic UMI counts were collapsed prior to generating the count matrix. To filter out ambient RNA barcodes, the number of UMIs per barcode were plotted with barcodes ordered by number of UMIs in descending order. The inflection point (or knee) in the plot was used as the threshold to separate barcodes originating from intact cells and barcodes originating from ambient RNA; all barcodes that fell below this threshold were removed. The remaining cells were further filtered for mitochondrial content (percent of UMI counts coming from mitochondrial genes) to remove low quality cells and total UMI counts to remove potential doublets with very large counts. The two sequencing batches were filtered separately. For sequencing batch 1 (D12, D14, D24, D26; see **Supplemental Table 1**), cells within the highest 5% of percent mitochondrial and total UMI counts distributions were removed. For sequencing batch 2 (D0, D90; see **Supplemental Table 1**), D90 cells had higher percentage mitochondrial UMIs than D0; this is consistent with previous studies showing an increase in mitochondrial content and number during cardiomyocyte maturation (Karbassi et al., 2020; Piquereau and Ventura-Clapier, 2018). D90 cells also had a decrease in total UMI counts compared to D0, which is consistent with other studies indicating a decrease after differentiation (Gulati et al., 2020). Due to these observations, we used the same filtering approach (removing the highest 5% of cells based on UMI and mitochondrial distributions) but applied it to each time point (D0, D12, D24 and D90) independently leading to different maximum cutoffs for each time point. For both sequencing batches, genes detected in less than 10 cells, and cells with less than 200 transcribed genes, were removed from the matrix prior to clustering and visualization.

### scRNA-seq clustering and visualization

Cells were clustered and visualized using the R (version 3.5.1) package Seurat (version 2.3.4) (Butler et al., 2018; R Core Team, 2018). Normalized transcript abundance for each gene was calculated by dividing counts by the total counts per cell, multiplying by a scaling factor (10,000) and log transforming the result using log1p (NormalizeData function in Seurat). For stem cell (D0) and protocol 1 samples (D12, D24, D90), highly variable genes to be used for dimensionality reduction and clustering were identified using FindVariableGenes with the following parameters: mean.function = ExpMean, dispersion.function = LogVMR, x.low.cutoff = 0.05, x.high.cutoff = 4, y.cutoff = 0.5 (these parameters were chosen to define outlier genes after manual inspection of mean expression vs. dispersion plot). Counts were scaled using ScaleData with default parameters without regressing out any variables. PCA was performed on the normalized and scaled matrix (with highly variable genes only) using RunPCA. Standard deviations of the principle components were plotted to determine the number of components to retain for clustering and visualization, and principle components above the inflection point in the plot were retained. Cells were clustered using the Jaccard-Louvain method, which is based on shared nearest neighbor modularity optimization (FindClusters function with resolution=c(0.3, 0.4, 0.5, 0.6, 0.8, 1) and the following standard parameters: algorithm=1 (original Louvain algorithm), modularity.fxn=1 (standard modularity function)). Uniform Manifold Approximation and Projection (UMAP) (McInnes et al., 2018) was used to visualize cells in a two-dimensional space (RunUMAP function with default parameters). Protocol 1 (D0 and small molecule D12, D24, D90; n = 11,619 cells) analysis is shown in Figure 1 and Figure 2; clustering with resolution 0.5 was used for visualization, differential expression and other downstream analyses. Clusters are labeled C0-C13 (Figure 1A, 1C). Pearson correlation was calculated for cardiac troponin T transcript and protein abundance between flow cytometry-based abundance (% cTnT positive cells, described in “Flow cytometry of scRNA-seq samples”) and scRNA-seq based abundance (% of *TNNT2* positive cells) (Figure 1E). Flow cytometry and scRNA-seq were performed on different cells obtained from the same differentiation sample (and therefore the same differentiation well). To compare differentiation protocols, cell lines, and differentiation experiments, Protocol 1 and 2 early and intermediate time point cells (D12, D14, D24, D26; n = 15,878 cells) were clustered together (clustering with resolution 0.8 is shown, and clusters are numbered C0-C16; **Supplementary Figure 1A-D**), and pairwise Spearman rank correlation coefficient was calculated (**Supplementary Figure 1E-H**). To more closely look at the relationship between D12 and D24 cardiomyocytes derived with protocol 1, cells from C0 (mostly D12) and C1 (mostly D24) (Figure 1A) were extracted, clustered, and visualized independently of all other cells first using the same workflow described above (clustering with resolution 0.5 is shown, and clusters are numbered i0-i5; **Supplementary Figure 2E-J**; n = 4,945 cells). Plots were created using R package ggplot2 (version 3.3.0; Wickham, 2016), and violin plots were created using R package scrattch.vis (version 0.0.210; Graybuck and Sedeno-Cortes, 2018).

### scRNA-seq differential expression analysis

Differentially expressed (DE) genes between clusters were identified by performing pairwise comparisons with edgeR (version 3.26.0) (version 3.26.0; McCarthy et al., 2012; Robinson et al., 2010). RNA composition normalization was performed with calcNormFactors, and negative binomial dispersions were estimated with estimateDisp. DE genes were identified by fitting gene-wise generalized linear models (glmFit; did not include intercept term in design) and performing a likelihood ratio test (glmLRT with contrasts). We retained genes with an absolute log2 fold change >= 1 and Benjamini-Hochberg adjusted p-value < 0.05). To remove potential false positives caused by dropouts in low depth data (i.e. gene expressed very highly in only a few cells within a cluster), we calculated the fraction of cells positive for each gene and retained only genes where the up-regulated group had at least 30% of cells positive for that gene. Heatmap of top differentially expressed genes (top 10 up and top 10 down regulated genes from each pairwise comparison by log2 fold change) (Figure 2A) was created using R package pheatmap (version 1.0.12; Kolde, 2019) and shows scaled transcript abundance (normalized counts for each gene were scaled and centered using Seurat’s ScaleData). For visualization purposes, maximum scaled transcript abundance value used in heatmap was +4, and values greater than that were set to 4. Genes in the heatmap were clustered using hierarchical clustering. Cells were grouped by cluster (shown in Figure 1A, 1C). All statistical calculations were performed in R 3.5.1, and plotting was performed using ggplot2 (version 3.3.0; Wickham, 2016).

### scRNA-seq feature selection analysis

In addition to differential expression analysis based on pairwise cluster comparisons, an orthogonal clustering-independent method was used for identifying genes that change between early and intermediate differentiation time points (D12 and D24). We fit generalized linear models with an elastic net penalty using the glmnet R package (version 2.0-13; Friedman et al., 2010; Zou and Hastie, 2005) alpha = 0.5 and a sequence of 100 values of lambda, the regularization parameter. The response variable was time point (D12 vs. D24). The input count matrix was filtered down to gene symbols that had a corresponding Entrez Gene ID, and cells were filtered to include only cardiomyocytes (cells positive for cardiomyocyte marker gene *TNNT2*). Cells were split into a training set (90% of cells; n = 4,283 D12 cells and n = 2,014 D24 cells) and a test set (10% of cells; n = 481 D12 cells and n = 227 D24 cells) and 1,000 bootstrap rounds (sampled 80% of cells without replacement at each round) were run with the training set to identify genes that had non-zero coefficients in all 1,000 rounds. Genes with non-zero coefficients in all 1,000 bootstrap rounds at lambda = 0.0494 were reported as selected (Figure 2B). The resulting list of 40 selected genes was used to predict the time point for each cell in the test set by fitting a binomial generalized linear model (glmnet with ridge penalty, alpha = 0, and lambda = 1×10^-6)^ with the training data set and then using the model to predict time point in the test data set. In addition to the 40 selected genes at lambda 0.0494, we fit models with gene sets selected at higher (more sparse; less genes) and lower (less sparse; more genes) values of lambda and calculated their accuracy in predicting time point in the test data set (Figure 2D). Accuracy was calculated using the confusionMatrix function from the R package caret (version 6.0-85; Kuhn, 2020) with positive = D12 time point. As a control for each selected gene set size, we took 100 random gene samples of the same size and calculated accuracy of predicting time point in the test data set, which was 0.68 D12 and 0.32 D24 cells. Accuracies are shown as box plots (outliers hidden) for random gene sets and as individual points for selected gene sets (Figure 2D).

### Replating cardiomyocytes for imaging and RNA FISH

All cardiomyocytes generated for RNA FISH (fluorescent *in situ* hybridization) and alpha-actinin-2-mEGFP imaging were differentiated using an optimized version of Protocol 1 in which Chiron and IWP2 were both used at 7.5 µM (refer to “Directed cardiomyocyte differentiation” section in Materials and Methods) (Roberts et al., 2019). After performing cardiomyocyte differentiation, cells were dissociated into single cells at D12 (described in “Single cell dissociation for scRNA-seq” section); 3 wells of a 6-well plate were dissociated into single cells, combined and used to seed multiple glass bottom multiwell plates. Flow cytometry was performed on a subset of cells for cardiac troponin T analysis as described in the “Flow cytometry of scRNA-seq samples” section above. Prior to cell plating, glass bottom multiwell plates (24-well, Cellvis P24-1.5H-N) were treated with 0.5 M glacial acetic acid (Fisher Scientific #BP1185-500) at RT for 20–60 min and washed once with sterile milliQ (MQ) water. Wells were then treated with 0.1% PEI (Sigma Aldrich #408727-100ML) solution in sterile MQ water for 16–72 h at 4°C, rinsed once with DPBS and once with sterile MQ water. Finally, wells were incubated overnight at 4°C with 25 µg/mL natural mouse laminin (Gibco #23017-015) diluted in sterile MQ water. Laminin solution was removed immediately prior to replating; cells were seeded at a density of 35,000 to 50,000 cells per well (24-well plate) in RPMI-1640 supplemented with B27 containing insulin, 1% P/S and 10 µM Rock Inhibitor. Media was changed after 24 hrs, and cells were maintained with RPMI-1640 supplemented with B27 containing insulin and 1% P/S every 2-3 days until fixation. Cells were fixed at time points indicated in text for RNA FISH and imaging of alpha-actinin-2-mEGFP structure. The D18 time point reflects single cell dissociation at D12 and a 6-day recovery period after replating, or additional maturation for D29-32 time points. Time points D18 and D32 were seeded from the same source population of D12 differentiated cardiomyocytes, replated together and maintained in parallel, then fixed at the appropriate time point. To fix cells, media was removed and wells were washed twice with RNAse-free PBS. Cells were fixed for 10 min at RT in a 4% paraformaldehyde solution (Electron Microscopy Sciences #15710), and then washed once more with RNAse-free PBS. PBS was removed and replaced with 70% ethanol; fixed plates were wrapped with parafilm and stored at −20°C until RNA hybridization was performed.

### RNA FISH using HCR v3.0

RNA FISH was performed using HCR v3.0, following the HCR v3.0 protocol for “Mammalian cells on a slide” with modifications to adapt for samples on glass bottom multiwell plates (Molecular Instruments, https://www.molecularinstruments.com/protocols). Wells were washed 4 times with 2X SSC (Invitrogen #15557-044), pre-hybridized in probe hybridization buffer (Molecular Instruments) for 30-60 min at 37°C, and hybridized overnight at 37°C with 1.2 pmol of each probe set mixture containing 400 U/mL RNAse inhibitor (Enzymatics #Y9240L). Custom probe sets were designed by Molecular Instruments (see **Supplemental Table 3**). After hybridization, wells were washed with probe wash buffer (Molecular Instruments) supplemented with 400 U/mL RNAse inhibitor at 37°C for 30 min and washed again 4 times with 2X SSC at RT. Amplification buffer (Molecular Instruments) was added and incubated for 30-60 min at RT. During this incubation, 18 pmol of hairpin amplifiers were heated to 95°C for 90 sec, protected from light and cooled, then combined and added into amplification buffer containing 400 U/mL RNAse inhibitor. Hairpin mixtures were incubated with the appropriate wells while protected from light for 45 min at RT for single molecule and diffraction-limited amplification. Excess hairpins were removed and wells were again washed with 2X SSC four times. Nuclei were labeled with 2 µg/mL DAPI in 2X SSC for 5 min, followed by additional washes with 2X SSC. Samples were stored protected from light at 4°C in 2X SSC with 400 U/mL RNAse inhibitor until imaging. Wash volumes used were 500 µL, and all other incubation steps used 250 µL per well, added using a multichannel pipette or P1000 in a 24 well plate format.

### Imaging replated cardiomyocytes

Imaging of AICS-0075 ACTN2-mEGFP cardiomyocytes (Figures 3-6**, Supplemental Figures 3, 5-6**) was performed on a 3i spinning-disk microscope with a 63x/1.2 NA W C-Apochromat Korr UV-vis IR objective (Zeiss) and a 0.83x tube lens adapter for a final magnification of 52.29x, a CSU-W1 spinning-disk head (Yokogawa) and a Prime BSI sCMOS Camera (Photometrics) (pixel size 0.120 µm in X-Y and 0.33 µm in Z). Imaging of unedited AICS-00 WTC cardiomyocytes for initial FISH target gene selection (Figure 2E-F, **Supplemental Figure 2M**) was performed on a Zeiss spinning-disk microscope with a 40x/1.2 NA W C-Apochromat Korr UV-vis infrared (IR) objective (Zeiss) and a 1.2x tube lens adapter for a final magnification of 48x, a CSU-X1 spinning-disk head (Yokogawa), and Orca Flash 4.0 camera (Hamamatsu) (pixel size 0.271µm in X-Y after 2×2 binning and 0.29 µm in Z). Standard laser lines (405, 488, 561, 640 nm), primary dichroic (RQFT 405, 488, 568, 647 nm) and the following Band Pass (BP) filter sets (Chroma) were used for fluorescent imaging: 450/50 nm for detection of DAPI, 525/50 nm for detection of mEGFP, 600/50 nm for detection of Alexa 546 dye and 690/50 nm for detection of Alexa 647 dye. For all live cell imaging (related to the “ML-based alpha-actinin-2 pattern classification” section below), cells were imaged in phenol red-free RPMI-1640 (Gibco #11835030) supplemented with B27 containing insulin media, within an incubation chamber maintaining 37°C and 5% CO_2_. In experiments using unedited AICS-00 WTC cardiomyocytes for initial FISH target gene selection, cell fixation occurred at either D18, or D32 ∓ 3 days as indicated in the text and relevant figures. In experiments using AICS-0075 ACTN2-mEGFP cardiomyocytes, cell fixation occurred at D18 or D32 as indicated. Brightfield images were acquired using an LED light source with peak emission of 740 nm with narrow range and a BP filter 706/95 nm for brightfield light collection.

### Manual cell annotations in Napari

Multi-channel Z-stacks were loaded into Napari (https://napari.org; napari contributors, 2019) and 2D single cell masks were generated by manually drawing cell boundaries in 2D while incorporating information from all channels collected during imaging (brightfield, two FISH probe channels, nuclei via DNA dye (DAPI), and alpha-actinin-2-mEGFP (structure) signal if present). Single cell masks in fields of view (FOVs) from RNA FISH experiments on unedited AICS-00 WTC cells (non-structure FISH) were generated by a single human expert. Single cell masks in FOVs from each alpha-actinin-2-mEGFP/FISH experiment well (structure/FISH) were randomly split into three groups and generated by one of three expert human annotators. Cell boundaries were manually drawn for cells that were mostly within the FOV, and low-confidence/high cell density regions with many overlapping cells were avoided. To speed up loading time for structure/FISH images in Napari, each image was down-sampled prior to manual annotation using the scale function in ImageJ, converting each 1776 x 1736 pixel x 50 planes Z-stack to an 888 x 868 pixels X 35 plane Z-stack using bilinear average interpolation. The 2D single cell masks were then rescaled to their original size prior to RNA FISH probe quantification. Non-structure FISH images were smaller in pixel dimensions and were not down-sampled. 2D single cell masks were used downstream to create single cell transcript abundance measurements, manual annotations of cell organization, single cell organizational scores, and integrated transcript abundance and organizational analysis in single cells (see “RNA spot segmentation and feature extraction”, “Expert annotation of cell organization”, “ML-based alpha-actinin-2 pattern classification”, and “FISH + alpha-actinin-2 assay data preparation” sections in Materials and Methods).

### DNA (Nuclear) segmentation

Nuclear segmentation in 2D using the DNA channel was performed using CellProfiler (version 3.1.8; McQuin C, 2018). See CellProfiler pipeline included in the corresponding github repository for details (Kamentsky et al., 2011). The DNA channel maximum intensity projection (MIP) was normalized with CellProfiler’s RescaleIntensity module from the 5^th^ percentile to 95^th^ percentile of the raw image. Nuclei were segmented using minimum cross entropy thresholding to define the probability distributions of foreground and background regions in an image using CellProfiler’s IdentifyPrimaryObjects module. Clumped objects were filtered by shape to identify nuclear objects in close proximity. Objects smaller than 5000 pixels (for images acquired on the 3i spinning-disk microscope; all *ACTN2*-mEGFP structure imaging, see “Imaging replated cardiomyocytes” section) or 500 pixels (for images acquired on the Zeiss spinning-disk microscope; all WTC non-structure imaging, see “Imaging replated cardiomyocytes” section) were considered debris and discarded. Nuclei were assigned to a cell if their centroids fell within the 2D segmented cell object. Unassigned nuclear objects were discarded and not used for further analysis.

### RNA spot segmentation and feature extraction

Channels with labelled RNA FISH transcripts were quantified using the Allen Cell Structure Segmenter (Allen Institute for Cell Science, 2020; Chen et al., 2018) by creating a transcript-specific segmentation workflow. MIP image intensities were normalized to fall between 3 standard deviations below the mean image intensity and 18 standard deviations above the mean intensity, and a Gaussian smoothing filter was applied to the images. A 2D spot filter algorithm was applied to the normalized images to segment the transcript signal. Each transcript signal was assumed to represent the location of the RNA species as a diffraction-limited spot. This filter takes into account both the radius of the dots and the filter response to generate the binary results. After visual inspection of images, we estimated the dot diameter to be 1-2 pixels and thus set the default value of the dot radius parameter to 1. Dot intensity levels varied by transcript, so we optimized the filter response parameter values individually for each RNA species to account for this variation as shown in the table below (probe set is indicated by gene name and corresponding amplifier wavelength):

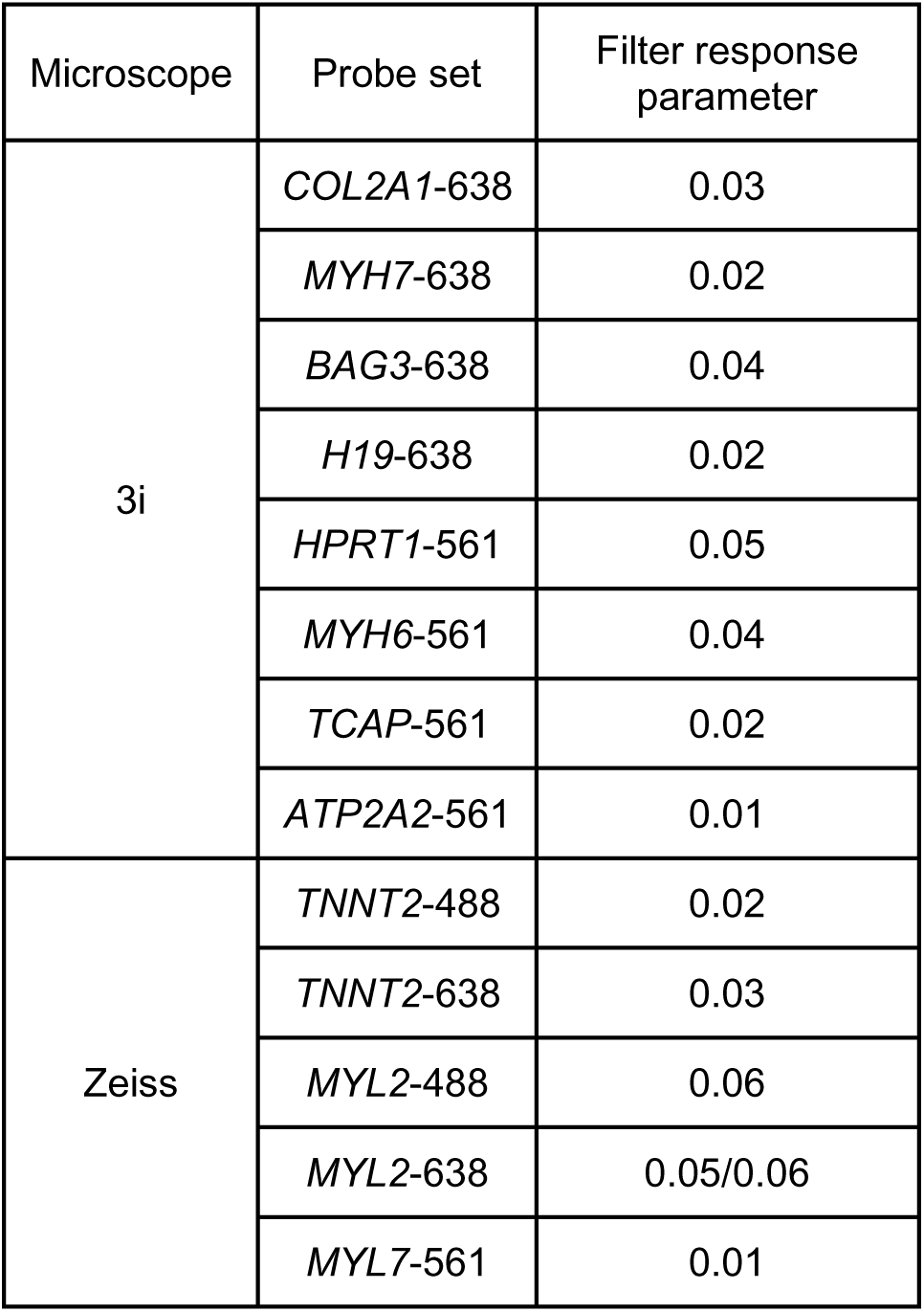
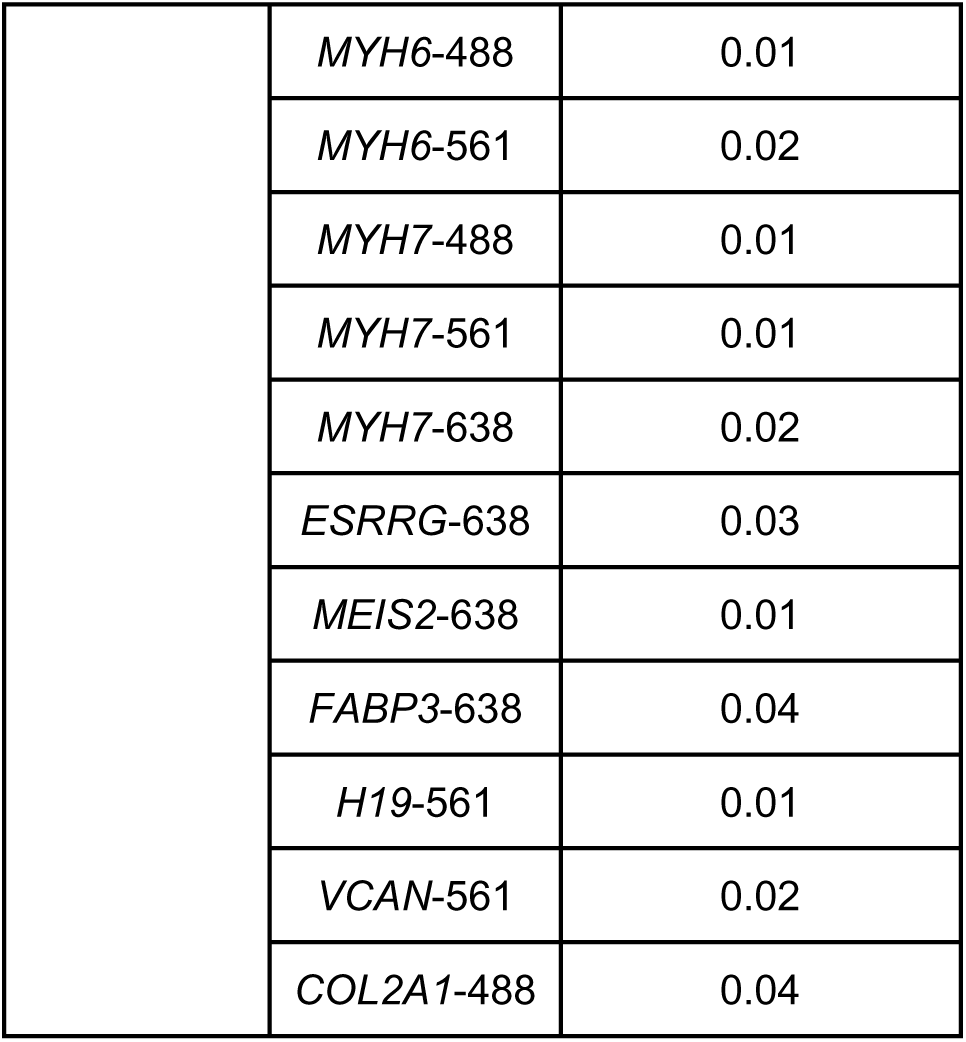

### Transcript localization

The localization and spatial distribution of transcripts in single cells were measured using three localization parameters for each transcript. First, the absolute distance between each segmented RNA species (see “RNA spot segmentation and feature extraction”) and the segmented nucleus (see “DNA (Nuclear) segmentation”) was calculated by computing the Euclidean distance between the center of mass of the transcript object and the center of mass of the segmented nucleus. Second, we calculated the Euclidean distance between the transcript and the center of mass of the cell. We then calculated the relative distance of a transcript from the nucleus vs. the cell boundary in single cells (Figure 6B, **Supplemental Figure 6C**) by taking the ratio of the nearest distance of the transcript to the nearest nuclear boundary (d_nucleus_), and the nearest distance of the transcript to the nearest cell boundary (d_cell_) using the single cell masks (see “Manual cell annotations in Napari”). Per cell reported values are the average relative distance of all segmented RNA spots for a given transcript (n = 1,287 cells for *MYH6* and *MYH7*, n = 1,190 cells for *COL2A1* and *HPRT1*, n = 1,156 cells for *H19* and *ATP2A2*, n = 1,142 cells for *BAG3* and *TCAP*).

Specific localization of *MYH6* and *MYH7* transcripts to sarcomeres, as identified by proximity to alpha-actinin-2 signal, was manually classified (n = 715 cells at D18 and n = 572 cells at D32, Figure 6C-D, **Supplemental Figure 6D-E**). If a gene did not have sufficiently dense transcript abundance to determine localization it was categorized as having “Low Expression”. If at least an estimated 30% of the transcripts co-localized with myofibrils, that cell was categorized as having sarcomeric transcript localization (see **Supplemental Figure 6A**). All other cells were categorized as having non-sarcomeric transcript localization. Manual scoring of *MYH7* transcript localization was further compared to the cell’s combined organizational score and regions with organized z-disks (Figure 6E, **Supplemental Figure 6F**, refer to “ML-based alpha-actinin-2 pattern classification”, “Linear model predicting expert organization scores from single cell metrics” and “Alpha-actinin-2 transcript localization enrichment scores” sections for details).

### Expert annotation of cell organization

Two experts manually scored each segmented cell for its overall alpha-actinin-2 organization. Each expert was presented with single cells cropped from a FOV (after manual annotation of single cell masks, see “Manual cell annotations in Napari” section for details); all cells were scored twice, once by each expert. Cells were scored as falling into one of five categories from 1-5; Figure 3B shows a schematic of the patterns that are representative for each class. Cells assigned a score of 1 had sparse, randomly organized puncta; cells assigned a score of 2 had denser puncta that were more organized; cells assigned a score of 3 exhibited a mix of puncta and other structures, including fibers and z-disks; cells assigned a score of 4 exhibited predominantly regular z-disks without a single clear axis of alignment; and cells assigned a score of 5 exhibited almost exclusively regular z-disks with a clear axis of alignment. Experts made their score based on the majority of the cell’s organization falling into one of these five categories. For example, a cell that had predominantly regular z-disks, but some puncta and fibers would still be categorized as a 4. Cells with approximately half regular structures and half irregular structures were nearly always categorized as a 3 (see **Supplemental Figure 3A** for additional examples). After all cells in the data set had been scored by both experts, annotations that differed by more than two units were curated; in general, these differences reflected cells with very low intensity alpha-actinin-2 signal. In these cases, experts established concordance after review. A total of 4,775 cells were scored (Figure 3C**, Supplemental Figure 3B**).

### ML-based alpha-actinin-2 pattern classification

A machine learning (ML) based alpha-actinin-2 pattern classifier was created based on images of live replated cardiomyocytes (AICS-0075 ACTN2-mEGFP) imaged on a 3i spinning disk microscope as described above in the “Imaging replated cardiomyocytes” section. This classifier was then validated for use in images of fixed alpha-actinin-2 images of the type used in this study (**Supplemental Figure 3I**). An expert manually annotated 3589 regions of 18 representative z-stacks of alpha-actinin-2-mEGFP into five classes according to their alpha-actinin-2 pattern: diffuse/messy, fibers, disorganized puncta, organized puncta, and organized z-disks. Fibers correspond to elongated thread-like patterns; diffuse/messy regions captured areas with many overlapping patterns or lacking discernible organization; disorganized puncta were assigned to small punctate structures with no obvious orientation; organized puncta were used to categorize regions with punctate structures that were organized along a discernible axis; and organized z-disks were used to capture well-aligned, stripe-like structures. We used Napari (https://napari.org; napari contributors, 2019) to click on and annotate pixel-based regions for each class; only the slice with the highest average intensity was used for manual annotation. We looked for regions in each image that clearly corresponded to each pattern category and captured the variation in these patterns. While we captured over 500 annotations for each pattern class, we did not fully annotate all regions of any image in the training set. Representative examples of these classes are shown in Figure 3E and **Supplemental Figure 3C-D**; this annotation was meant to capture the variety of patterns observed in the data set.

To identify background regions, we used the `filament3d` Jupyter notebook workflow template from the Allen Cell Structure Segmenter to segment the images in 3D (Chen et al., 2018), and then converted the 3D stack into a 2D image by keeping only the slice with the highest average intensity. Next we tiled the image with tiles of size 64×64 pixels and computed the total number of foreground pixels inside each tile; tiles with less than 64 foreground pixels were flagged for the next step. We calculated the distance between every pixel in the flagged tiles to the closest foreground pixel. If that distance was greater than 64, we classified that pixel as background.

Each non-background annotation was stored as an (X,Y) coordinate and used as training data for a deep-learning-based classifier model. Training data was drawn from live cell imaging of replated alpha-actinin-2-mEGFP (AICS-0075 ACTN2-mEGFP) cardiomyocytes. This data was split into train/test data sets, with 80% of the data being used for training and 20% held out as a testing set. We used a ResNet18 convolutional neural network pre-trained on the ImageNet data set as our classifier. All layers of this network were fine-tuned, and the last layer was trained from scratch on our training set. We used patches of 96 x 96 pixels centered around each annotation, upsampled to 256 x 256 pixels through linear interpolation, as input to the network. The network was trained to predict the class of the center pixel of the patch. We trained the model for 750 epochs and we applied the trained model to the test set, where the model performed with 83% accuracy (see **Supplemental Figure 3E**). We used a separate data set of alpha-actinin-2-mEGFP cardiomyocytes that was imaged both in live and fixed conditions to determine how the fixation process impacts the classifier performance. Because the alpha-actinin-2 signal from live cell imaging was significantly brighter than the signal in the matched fixed images, the histograms of the fixed images were matched to the average histogram of the live cell training data set prior to applying the model. **Supplemental Figure 3I** shows that there are only minor changes in the overall prediction suggesting that the fixation process has negligible impact for use of this model. The intensities for the FISH data set were not normalized to live cell imaging intensities as their signal was adequate for classifier performance. The ML model was used to make pixel-wise predictions of the alpha-actinin-2-mEGFP pattern classes on the highest intensity slice of all fields of views used in this paper. For these predictions, each image patch was evaluated by the model 4 times: once in an unaltered state, and three times with the patch undergoing randomized mirroring and rotation. The class probabilities from each round were averaged together for the final classification. The resulting classification maps were masked based on the single cell masks (see “Manual cell annotations in Napari” section) to calculate and output the fraction of cell area covered by each alpha-actinin-2-mEGFP pattern class on a per cell basis.

### Quantification of global sarcomere alignment

To quantify global structural alignment, we implemented a method described in (Sutcliffe et al., 2018) which uses the correlation of a gray-level co-occurrence matrix to quantify patterns of sarcomeric organization. In this approach, the element (*i*, *j*) of the co-occurrence matrix of an image *I* of dimension *N* × *M* is defined as

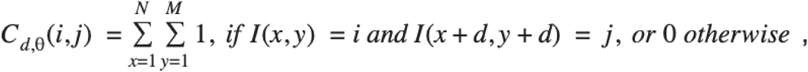

and it represents the number of times that grey-level *j* occurs at an offset distance *d* and at an angle *θ* from grey-level *i*. The correlation of the co-occurrence matrix measures the joint probability of pairs of gray-levels and it is defined as

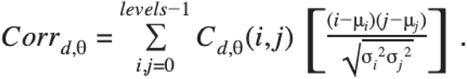

The correlation function displays a fast exponential-type decay in cells with no regular patterns. The correlation function oscillates when regular patterns with no preferential direction of alignment are present (see **Supplemental Figure 3J**). A mix of exponential-decay and oscillatory behavior is observed in cases where a preferential direction of alignment is present (**Supplemental Figure 3K**). In (Sutcliffe et al., 2018) the authors suggest using the height of the highest peak as a score for sarcomere organization. To detect the highest peak, we first fit the exponential decay component of the correlation curves as shown by the dashed lines top graph of **Supplemental Figures 3J-K**. Next, we subtracted the fitting curves from the raw data to produce the curves shown in the middle plot of **Supplemental Figures 3J-K**. The height of the highest peak is easily identified in the curves and shown by the horizontal dashed line, and green arrow labeled “peak height”. This value was used as a metric for sarcomere spacing.

We also used the maximum coefficient of variation of the correlation signatures as a metric for global sarcomere alignment. This metric is particularly useful for distinguishing between aligned and non-aligned fibers. In the case of well aligned fibers, we see heterogeneity in the Haralick decay profiles, while for poorly aligned fibers, the decay profiles are homogeneous. This difference is effectively captured by the coefficient of variation, even though the curves lack a clear peak. The maximum coefficient of variation is defined as:

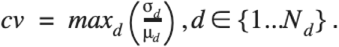

The mean response *μd* for a given distance *d* is defined as

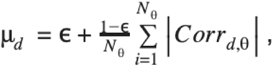

and the standard deviation is given by

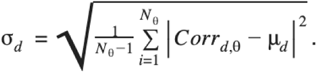

The variable *ε* is a numeric regularization factor chosen to be 0.1. The total number of values of *d* was *N_d_* = 32 and the total number of different angles was *N_θ_* = 16 for all experiments. Prior to co-occurrence matrix calculation, every cell image was converted to 3-bit format (*levels* = 8) with the same number of pixels in each level. Background was not included in the co-occurrence matrix calculation. The bottom right corner of **Supplemental Figures 3J-K** show the coefficient of variation, mean response and standard deviation in cells where the maximum coefficient of variation is low and high, respectively.

### FISH + alpha-actinin-2 assay data preparation

For all analysis of the relationship between transcript abundance (RNA FISH) and alpha-actinin-2 structural organization, we used a data set of 477 five-channel 3D FOVs, projected into 2D by taking the maximum intensity along the z-axis. From these 2D fields, we manually segmented 4,775 single cell masks using Napari (see “Manual cell annotations in Napari”). For each single cell mask we then had five channels of data: two RNA FISH probe channels, one channel of fluorescently tagged alpha-actinin-2 protein, a brightfield channel, and a fluorescent nuclear dye channel (DAPI). We assayed four pairs of FISH probes: *HPRT1* and *COL2A1* (n = 1,190 cells), *MYH6* and *MYH7* (n = 1,287 cells), *TCAP* and *BAG3* (n = 1,142 cells), and *ATP2A2* and *H19* (n = 1,156 cells) at two time points (D18 and D32); all other channels were identical throughout the experiment. In addition to the raw image channels, segmentations were performed for all channels other than the brightfield. Both the raw and normalized 2D images and segmentations are available in the data supplement.

From these five channel images, we constructed a variety of quantitative metrics, which are available in tabular form in the supplemental data. The quantitative metrics fall into four categories:

1. Cell features: the segmented images were processed by CellProfiler to generate cell area and aspect ratio measurements for each cell. An exhaustive list of single cell features was generated and is included in the corresponding quilt data package.
2. Expert scores of sarcomeric organization: the structural maturity of the cells was assessed by two human experts, on an ordinal scale of 1–5. See “Expert annotation of cell organization” section.
3. Local organization: all pixels in individual cells were classified according to their alpha-actinin-2 pattern: diffuse/messy, fibers, disorganized puncta, organized puncta organized z-disks or background. These scores were aggregated at a single cell level as six numbers that represent the fraction of the cell area covered by each of these classes. See “ML-based alpha-actinin-2 pattern classification” section.
4. Global alignment: three metrics for global sarcomere alignment was calculated for each cell. See “Quantification of global sarcomere alignment” section.

We focus in on a smaller sub-list of 11 simple features for our modeling: two features that describe the cell’s gross morphology (cell area and cell aspect ratio), six features that describe the cell’s local structure (described in the “ML-based *ACTN2* pattern classification” section), and three features describing the cell’s global sarcomeric alignment (described in the “Quantification of global sarcomere alignment” section.).

All data used to generate figures in this manuscript is available at: https://open.quiltdata.com/b/allencell/tree/aics/integrated_transcriptomics_structural_organization_hipsc_cm

### Linear model predicting expert organization scores from single cell metrics

The “combined organizational score” used for further analysis in Figure 4 **and** Figure 5 was constructed using ordinary least squares regression, using the eleven cell features in Figure 4B to predict the human expert annotation scores, which are integer values from one to five. In order to maintain predictive fidelity to biologically important but low abundance extremal scores, the regression was fit using sample weights in inverse proportion to the abundance of the sample’s expert annotation score. The regression also included a minute ridge penalty for numerical stability. All features were standardized to zero mean and unit variance before fitting the model, in order to more reasonably assess relative importance of the features using the coefficients of the fitted model. Lastly, an offset term was calculated for the regression. All details are available in our github repository (https://github.com/AllenCellModeling/fish_morphology_code).

Once fit to the entire data set of 4,775 cells, the model was then applied to the data set to generate the “combined organizational score” for each cell as a linear combination of the input features, using the regression coefficients (and offset) of the linear model as weights. Because the number of features is so small compared to the number of data points, overfitting concerns are minimal. To assess the predictive capacity of these features on the relative expression of *MYH6* and *MYH7* (see “Differential expression between *MYH6* and *MYH7*” section below for details), a similar linear model was also fit regressing this quantity, using the same features, normalization, and regularization. The predicted and actual values of the *MYH7-MYH6* relative expression are shown in Figure 5F.

### Bootstrapped confidence intervals

Confidence intervals are calculated using bootstrap resampling (Efron, 1979; Wasserman, 2006). The data set is resampled (with replacement) 1,000 times, and the statistical operation of interest (Spearman correlation, linear regression, etc.) is computed on each bootstrap sample. The values for that statistic are then ranked, and the values at the 2.5 and 97.5 percentiles are reported as the lower and upper bounds of the 95% confidence interval.

### Probability densities

The marginal density histograms in Figure 2C, Figure 4E, F**, Supplemental Figure 4C**, Figure 5C, are normalized such that the area under each curve integrates to one, i.e. they are probability densities.

### Correlation analysis

For variables that are either ordinal or continuous, correlations are assessed via Spearman rank-correlation (Spearman, 1904), a robust correlation estimator that quantifies the monotonicity of the relationship between two variables. We treated our time point data as a binary categorical variable rather than as an ordinal variable; thus, differences between other continuous and ordinal variables as a function of day were assessed using a Mann-Whitney U-test (Mann and Whitney, 1947). This test is a non-parametric quantification of how likely it is that samples were drawn from the same distribution. Data tabulation and statistical calculations were performed in Python 3.7 using the Numpy 1.18.2, Scipy 1.4.1, Pandas 1.0.3, scikit-learn 0.22.2, AnnData 0.7.1, and Pingouin 0.3.3 libraries. Plotting was performed using the Altair 4.0.1 and Seaborn 0.10.0 visualization libraries (McKinney and others, 2010; Oliphant, 2006; Pedregosa et al., 2011; Vallat, 2018; Van Rossum and Drake Jr, 2009; VanderPlas et al., 2018; Virtanen et al., 2020; Waskom et al., 2020; Wolf et al., 2018). Data organization and distribution was performed using quilt3distribute 0.1.3 (Brown, 2019).

### Differential expression between MYH6 and MYH7

In the subset of cells (n = 1,287) where *MYH6* and *MYH7* were jointly probed via RNA FISH, an aggregate per-cell score was constructed to indicate the relative expression of *MYH6* vs *MYH7*, as follows. First, the segmented FISH spot counts for each gene in each cell (see “RNA spot segmentation and feature extraction” section) were normalized by the cell area to create a count density for each gene in each cell: *d_i_* ^*g*^ = *c_i_* ^*g*^/*a_i_*, where *c_i_* ^*g*^ is the count of gene *g* in cell *i*, *a_i_* is the area of cell *i*, and *d_i_* ^*g*^ is the density of gene *g* in cell *i*. Densities of each gene were then normalized by the median expression of each gene to account for the different scales of their respective expression densities, creating a normalized density metric: *n_i_* ^*g*^ = *d_i_* ^*g*^/*Mg* where *Mg* = *Mediani*(*di* ^*g*^) is the median expression density of gene *g* over all cells *i* in which it was probed. The relative expression density of *MYH7* to *MYH6* in each cell is then calculated as *r_i_* = (*n_i_*^*MYH*7^ − *n_i_*^*MYH*6^)/(*n_i_*^*MYH*7^ + *n_i_*^*MYH*6^).

### Alpha-actinin-2 transcript localization enrichment scores

We also wanted to develop a quantitative metric for the enrichment of transcripts to specific classes of alpha-actinin-2 organization (**Supplemental Figure 6F**). The y-axis of plots shown in **Supplemental Figure 6F** capture the *enrichment* of transcripts to specific regions of the cells, a metric which accounts for the size of the regions by assuming a null distribution of transcripts that are of uniform random distribution.

Let *n_i_* be the total number of *MYH7* fish spots in cell *i*, and *n_i_*^*k*^ the number of fish spots in cell *i* that are also in in the area of the cell labeled as category *k*. To construct the enrichment score for category*k* in cell *i*, we compute the fraction of transcript spots in the areas labeled as category *k* of cell *i*: *f_i_*^*k*^ = *n_i_*^*k*^/*n_i_*. We also compute the fraction of spots we would expect in that area at random, which is just the fraction of the cell area labeled as category *k*: *r_i_*^*k*^ = *a_i_*^*k*^/*a_i_*, where *a_i_* is the area of cell *i*, and *a_i_* ^*k*^ is the area of that cell labeled as category *k*. To compute the enrichment score *e_i_*^*k*^ we then subtract the latter correction from the former quantity: *e_i_*^*k*^ = *f*_i_^*k*^−*r*_i_^*k*^ = *n_i_*^*k*^/*n* − *a_i_^k^*/*a_i_*

## QUANTIFICATION AND STATISTICAL ANALYSIS

Details of specific statistical analyses for each section, sample sizes, and statistical tests used are given in the Materials and Methods and in the corresponding figure legends.

