## Supplementary figures and images for "Cell states beyond transcriptomics: integrating structural organization and gene expression in hiPSC-derived cardiomyocytes"

### Supplemental Figures

Supp. Figure 1

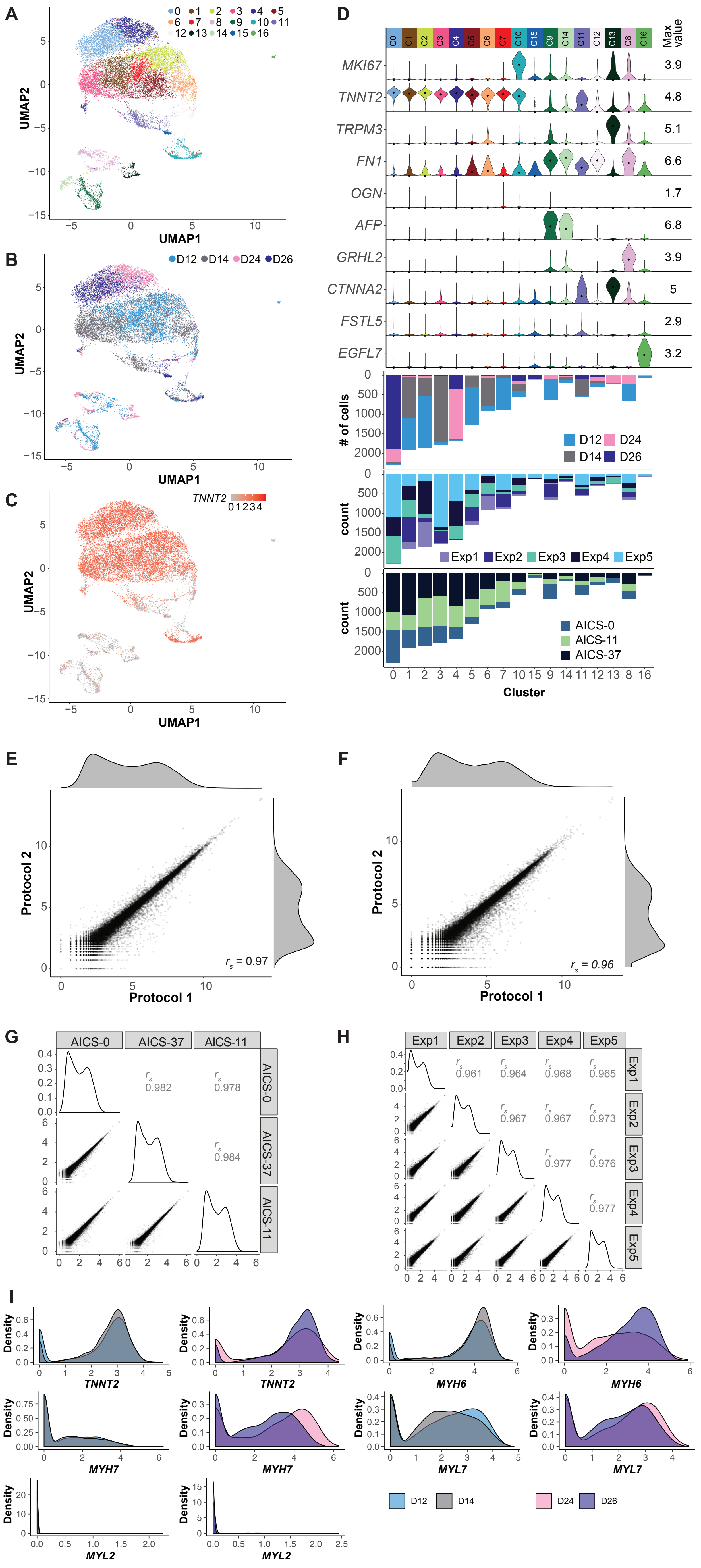

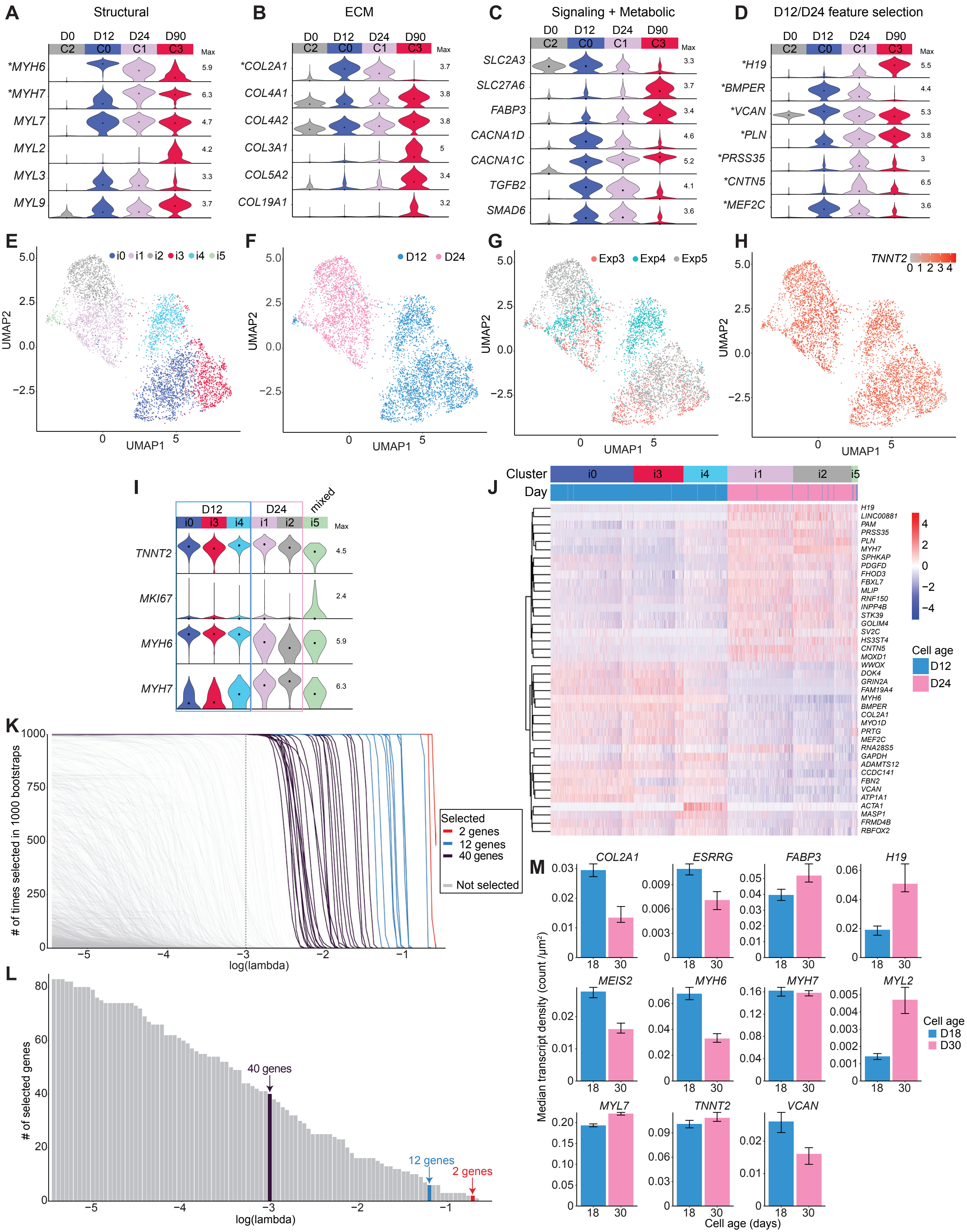

Supplemental Figure 3

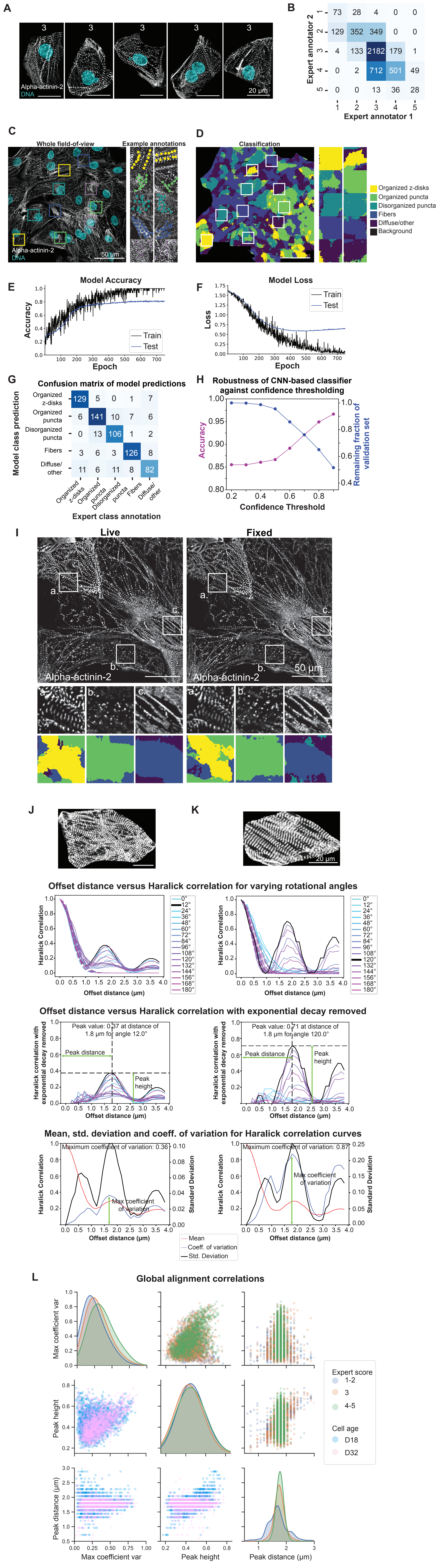

Supplemental Figure 4

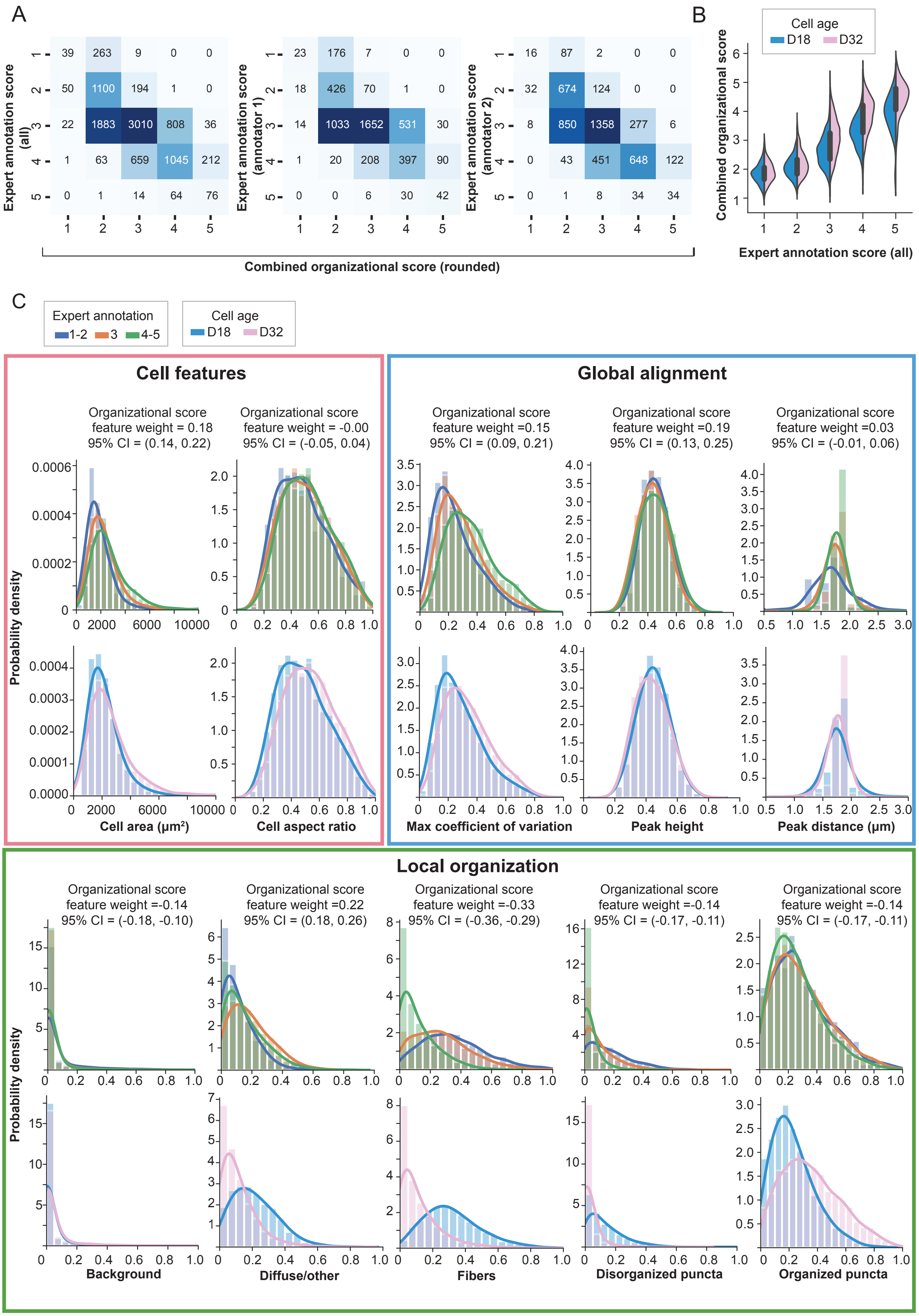

Supplementary Figure 5

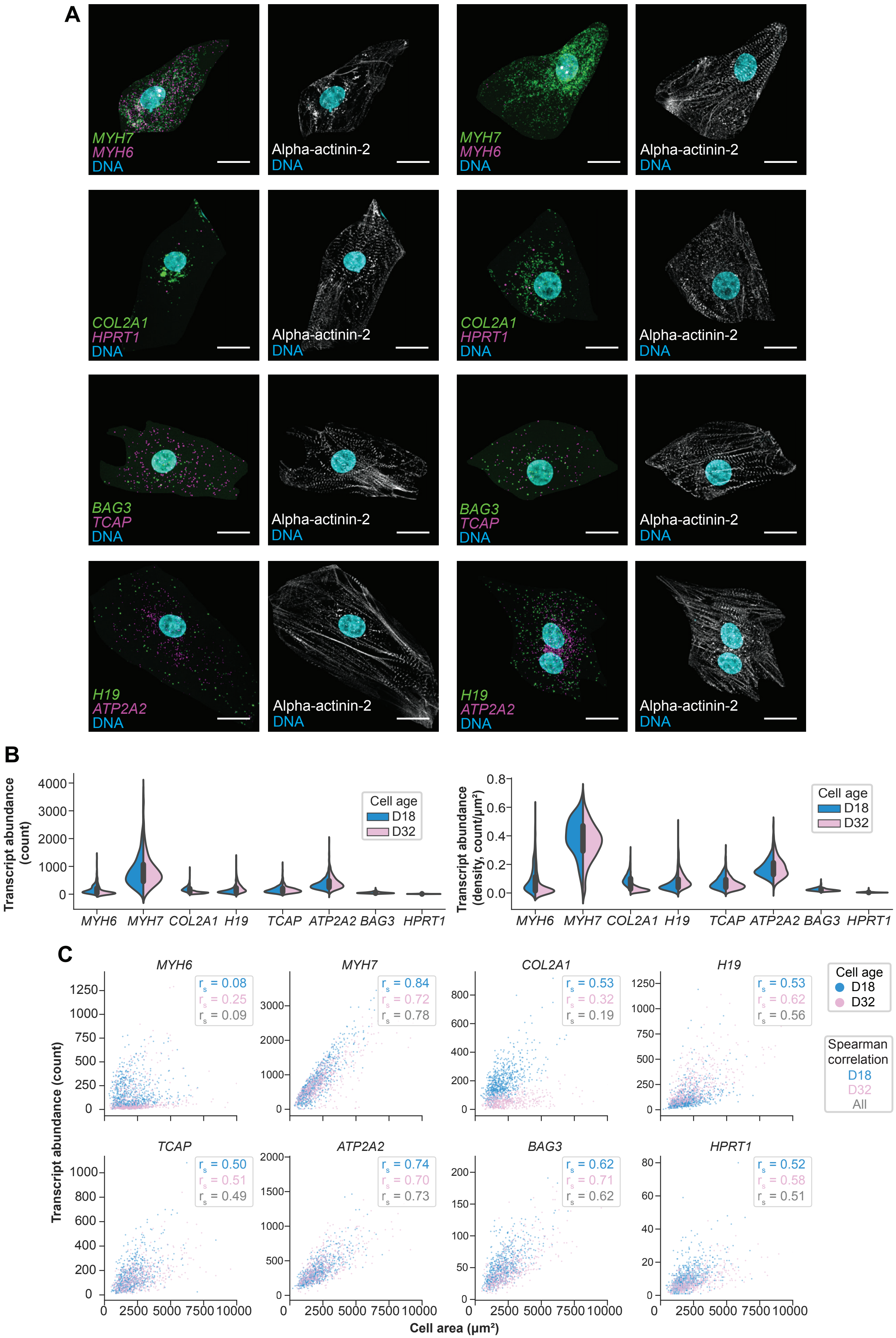

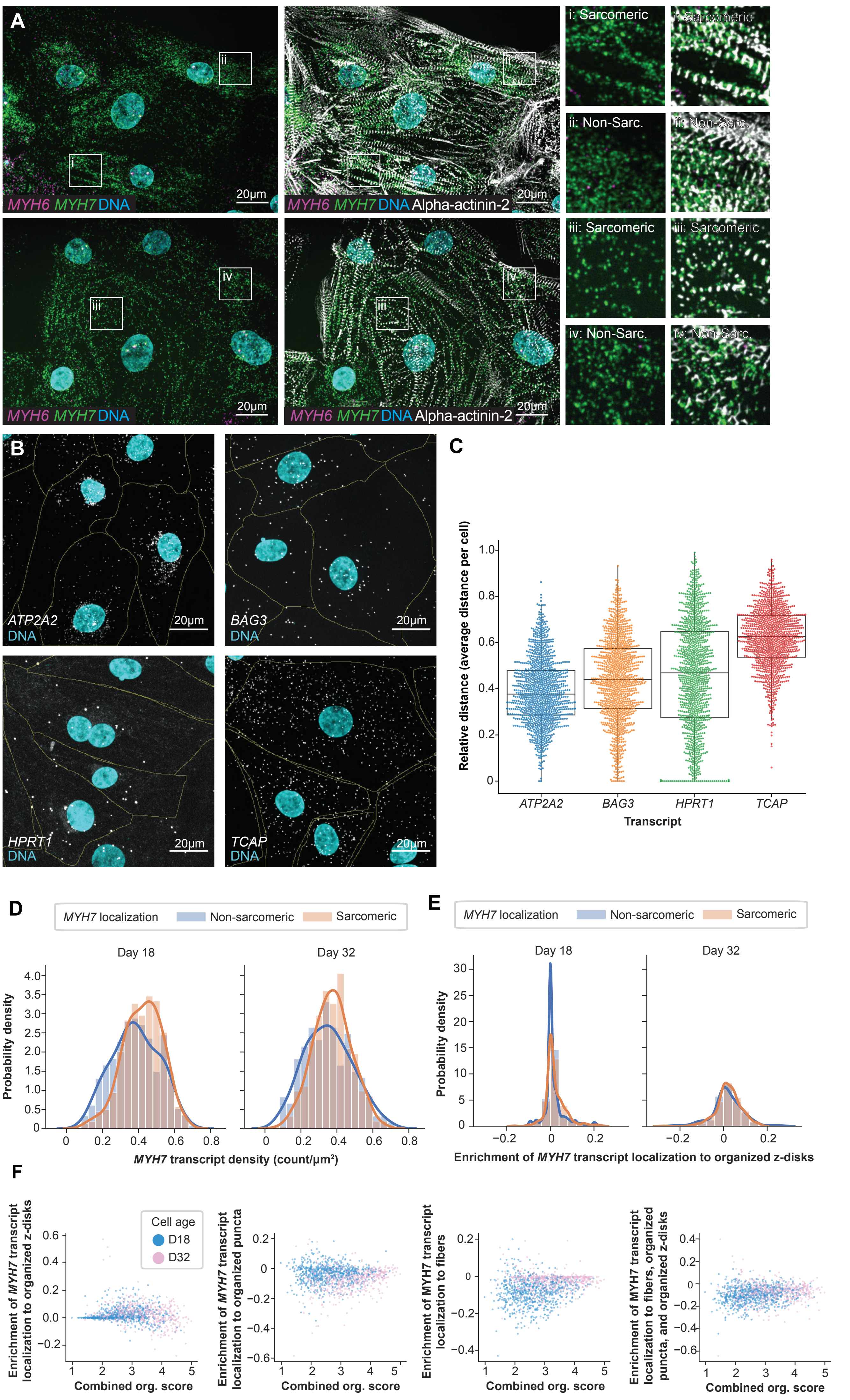
